# Genetically encoded biosensor for fluorescence lifetime imaging of PTEN dynamics in the intact brain

**DOI:** 10.1101/2024.10.04.616644

**Authors:** Tomer Kagan, Matan Gabay, Yossi Levi, Sharbel Eid, Nikol Malchenko, Maya Maman, Anat Nitzan, Luca Ravotto, Ronen Zaidel-Bar, Maayan Gal, Tal Laviv

## Abstract

The phosphatase and tensin homolog (PTEN) is a vital signaling protein which maintains an inhibitory brake that is critical for cellular metabolism, proliferation, and growth. The importance of PTEN signaling is evident from the broad spectrum of human pathologies associated with its loss of function. Moreover, loss or gain of PTEN function in animal models leads to aberrant cellular morphology, function, and metabolic regulation. However, despite the important role of PTEN signaling, there is currently no method to dynamically monitor its activity with cellular specificity within intact biological systems. Here, we describe the development of a novel PTEN biosensor, optimized for two-photon fluorescence lifetime imaging microscopy (2pFLIM). This biosensor is designed to measure PTEN activity within intact cells, tissues, and organisms. Our approach is based on monitoring FRET-dependent changes in PTEN conformation, which serves as a proxy for the activity state in living cells. We identify a point mutation that allow us to express this biosensor with minimal interference to endogenous PTEN signaling and cellular function. We demonstrate the utility of imaging PTEN signaling in cell lines, developing *C. elegans,* and in the living mouse brain. To complement this approach, we developed a red-shifted PTEN sensor variant that permits simultaneous imaging with GFP-based sensors. Finally, we use in vivo PTEN imaging in the mouse brain to identify cell-type specific dynamics of PTEN activity in excitatory and inhibitory cortical cells. In summary, our approach enables dynamic imaging of PTEN activity in vivo with unprecedented spatial and temporal resolution.

## Introduction

During development, cellular function and structure are regulated by key signaling pathways. As a corollary, developmental disorders and pathologies can be associated with specific impairments in these key regulatory genes. One striking example is the phosphatase and tensin homolog (PTEN) tumor suppressor gene ^1^. PTEN is a lipid and protein phosphatase that regulates cell growth, proliferation, and survival by dephosphorylating phosphatidylinositol 3,4,5-trisphosphate (PIP3), and thereby inhibiting the downstream activation of serine/threonine kinase (AKT) ^2^. PTEN signaling is regulated by several cellular pathways, such as growth factor receptors ^3^ and Rho family proteins ^4^. In addition, PTEN signaling is differentially regulated in distinct subcellular compartments, such as the mitochondria ^5^ and the nucleus ^6,7^. In the brain, PTEN signaling is vital for coordinating neuronal and glial development ^8–10^, synaptic plasticity ^11–15^, axonal growth and guidance ^16,17^, and cell survival ^2,18,19^. Dysregulation of PTEN has been closely associated with systemic neoplasia ^10,20,21^ and neurological disorders, including autism spectrum disorders (ASD) ^22–24^, PTEN hamartoma tumor syndrome (PHTS) ^25^, epilepsy ^26^, and macrocephaly ^23,24^. Although the vital and conserved roles of the protein across tissues and species has made PTEN the subject of extensive study, current methodologies for assessing PTEN function are limited to ex-vivo downstream biochemical assays, or to manipulations inducing loss or gain of function of PTEN. PTEN loss of function (LOF) in the nervous system leads to hypertrophy in neurons and astrocytes ^9,27^, as well as increased synaptic density and hyperconnectivity ^15,28,29^. Accordingly, systemic PTEN LOF results in ASD like behavioral features and severe neurodevelopmental pathology ^8,9,18,30^. Conversely, PTEN over-expression (OE) results in brain microcephaly, decreased synaptic density and neuronal dysfunction ^14,31^. Interestingly, systemic PTEN OE increases longevity, metabolic functions and oncogenic resistance ^32,33^. Endogenous regulation of PTEN signaling therefore constitutes a vital mediator of brain development and function. However, despite intense study, the precise dynamics of PTEN signaling activity in intact cells, tissues and organisms remain unknown. Previous studies that conditionally deleted PTEN in specific cell types have suggested that cell-specific PTEN function is critical for brain development ^18^. However, since genetic manipulations of PTEN lead to severe cellular malfunction, it remains unclear whether PTEN exerts cell-type specific spatial and temporal signaling activity within a physiological context.

PTEN was previously shown to undergo a conformational structural rearrangement upon activation, transitioning from a closed inactive compact conformation to active and open relaxed form ^34,35^. This characteristic was previously utilized to engineer a bioluminescence resonance energy transfer (BRET) based approach to assess PTEN activity in living cells ^36^. Unfortunately, this method could not achieve cellular resolution and is difficult to implement within an intact complex cellular environment, such as the brain. An Ideal experimental approach would allow monitoring of cell-type specific PTEN activity in vivo with subcellular resolution and temporal precision, and with minimal perturbation to the endogenous signaling.

Here, we describe a PTEN biosensor engineered to monitor conformational changes by FRET, as a proxy for PTEN activity levels. We used two-photon fluorescence lifetime imaging microscopy (2pFLIM) as a robust readout for PTEN activity. This approach was validated by pharmacological and genetic perturbations, thereby demonstrating the ability to monitor the state of PTEN activity in living cells. Moreover, we have used this approach to reveal how various pathogenic LOF point mutations in PTEN lead to structural destabilization, which were as evidenced by PTEN conformation. We then screened and identified an optimized biosensor variant containing a single point mutation. This variant, which we named G-PTEN (for GFP based PTEN sensor), maintains conformational flexibility to serve as a sensitive proxy of PTEN activity but exerts minimal perturbation on endogenous PTEN signaling. We then integrated G-PTEN in the model organism *C. elegans* and monitored developmental changes in PTEN activity state, and their dependence on the longevity related *daf-2* pathway. In addition, we utilized the PTEN biosensor for in vivo imaging in the intact mouse brain. The use of 2pFLIM in vivo, allowed us to monitor the PTEN activity state with cellular and subcellular precision, and to demonstrate how genetic perturbations within the PTEN signaling pathway led to subcellular alterations in PTEN signaling, which correspond to structural synaptic and cellular deficits. Finally, we engineered a red-shifted variant, R-PTEN, which allows simultaneous imaging of PTEN signaling and neuronal activity with a GFP based calcium indicator. The combination of spectral variants of the PTEN sensor enables PTEN activity to be monitored simultaneously in excitatory and inhibitory cells in vivo. This allowed us to reveal previously unknown distinct cell-type specific experience dependent changes in PTEN signaling.

Altogether, our novel biosensor combined with 2pFLIM is ideally suited for monitoring PTEN signaling within living cells, intact tissues, and whole organisms and can be used to identify precise cell-type specific physiological functions.

## Results

### Development of a new PTEN FRET/FLIM biosensor

We set out to develop a biosensor for monitoring PTEN activity in living cells. Our design is based on changes in the conformation of PTEN **(Fig. 1a**), which was shown to undergo a transition from a closed to an open state upon activation ^34,35^. Our goals were to design a cell specific reporter of PTEN activity dynamics, which would allow robust readout with subcellular resolution within complex innate tissue, such as the brain. We hypothesized that labeling the N and C termini of PTEN with donor and acceptor fluorescent proteins would enable FRET detected changes in PTEN conformation to serve as a proxy of its activity. This approach was previously used to engineer a BRET based PTEN sensor ^36^. We employed two-photon Fluorescence lifetime imaging (2pFLIM), a robust and quantitative FRET readout enabling monitoring of biosensor activity within intact complex tissue, such as the living mouse brain over extended time periods ^37–39^. We engineered PTEN with monomeric enhanced GFP (mEGFP) as a FRET donor, and sREACh, a dark YFP variant, as an acceptor. This combination allows to measure FRET/FLIM dynamics reflected by changes in GFP lifetime ^40,41^ and permits simultaneous imaging with other red-shifted biosensors ^37,42^. The mEGFP-PTEN constructs were expressed in HEK293 cells and engineered to optimize the linker regions between PTEN and the donor and acceptor fluorescent proteins (**Extended fig. 1**). This design was validated by mutating four phosphorylation sites at the C terminus of PTEN (Ser380, Thr382, Thr383, and Ser385) to alanine (4A). These mutations were previously shown to cause a constitutively active and open form of PTEN, due to loss of phosphorylation ^34^. We reasoned that higher dynamic range of the sensor would be reflected by larger FLIM differences between WT and 4A variants (**Extended fig. 1b-c)**. These efforts led to a sensor design with truncated linkers, which possesses high basal FRET level mirrored by low fluorescence lifetime, as compared to donor only tagged PTEN (**Fig. 1b-d**, 2.20 ± 0.003 ns for donor and acceptor, 2.69 ± 0.001 ns for donor only). The FRET values for the selected 4A mutant variant were dramatically reduced (**Fig. 1b-d**, 2.53 ± 0.002 ns), which confirms that the high basal levels of FRET of the normal biosensor are due to endogenous regulation by phosphorylation. As the next step, we assessed the ability of our biosensor to detect dynamic changes in PTEN activity over time. For this purpose, we used the tetrabromobenzotriazole (TBB), an inhibitor of casein Kinase 2 (CK2), which is the main kinase that phosphorylates and thereby inhibits PTEN activity ^43^. Application of TBB, resulted in a long-lasting increase of fluorescence lifetime, reflecting increased PTEN activity (**Fig. 1e-g**, 0.23 ± 0.003 ns, **Supplementary movie 1**). In contrast, TBB treatment of cells expressing the 4A mutant PTEN sensor, caused minimal changes in FRET (**Fig. 1e-g**, −0.06 ± 0.003 ns), which confirms the association with phosphorylation induced activity. To determine whether the PTEN biosensor is amenable to bidirectional regulation, we applied epidermal growth factor (EGF), which negatively regulates PTEN activity ^44^. This resulted in a significant decrease of PTEN activity (**Fig. 1h-i**, Control-2.23 ± 0.003 ns, EGF-2.17 ± 0.003 ns). The relatively small change in lifetime and decrease in PTEN activity observed may be due to the low basal level of PTEN activity in proliferating cells such as HEK cells, since PTEN signaling is pro-apoptotic. In order to address this issue and examine the reversibility of PTEN conformational dynamics, we first applied TBB to increase PTEN activity followed by EGF application. This manipulation induced significant inhibition of PTEN, which was not due to washout of TBB (**Fig. 1j-k**). Finally, we examined whether the PTEN biosensor could detect regulation of upstream signaling pathways, which were previously associated with PTEN. For this purpose, we overexpressed variants of RhoA kinase, which is associated with cellular proliferation and growth and has been reported to regulate upstream PTEN signaling ^4^. Overexpression of the constitutively active form of RhoA led to a significant increase in lifetime, which corresponds to PTEN activation, while a dominant negative RhoA mutant did not alter the basal PTEN activity state (**Fig. 1l**). We can therefore conclude that our experimental approach enables us to monitor PTEN conformation using FRET/FLIM, as a reliable proxy of PTEN activity in living cells.

**Figure 1.**
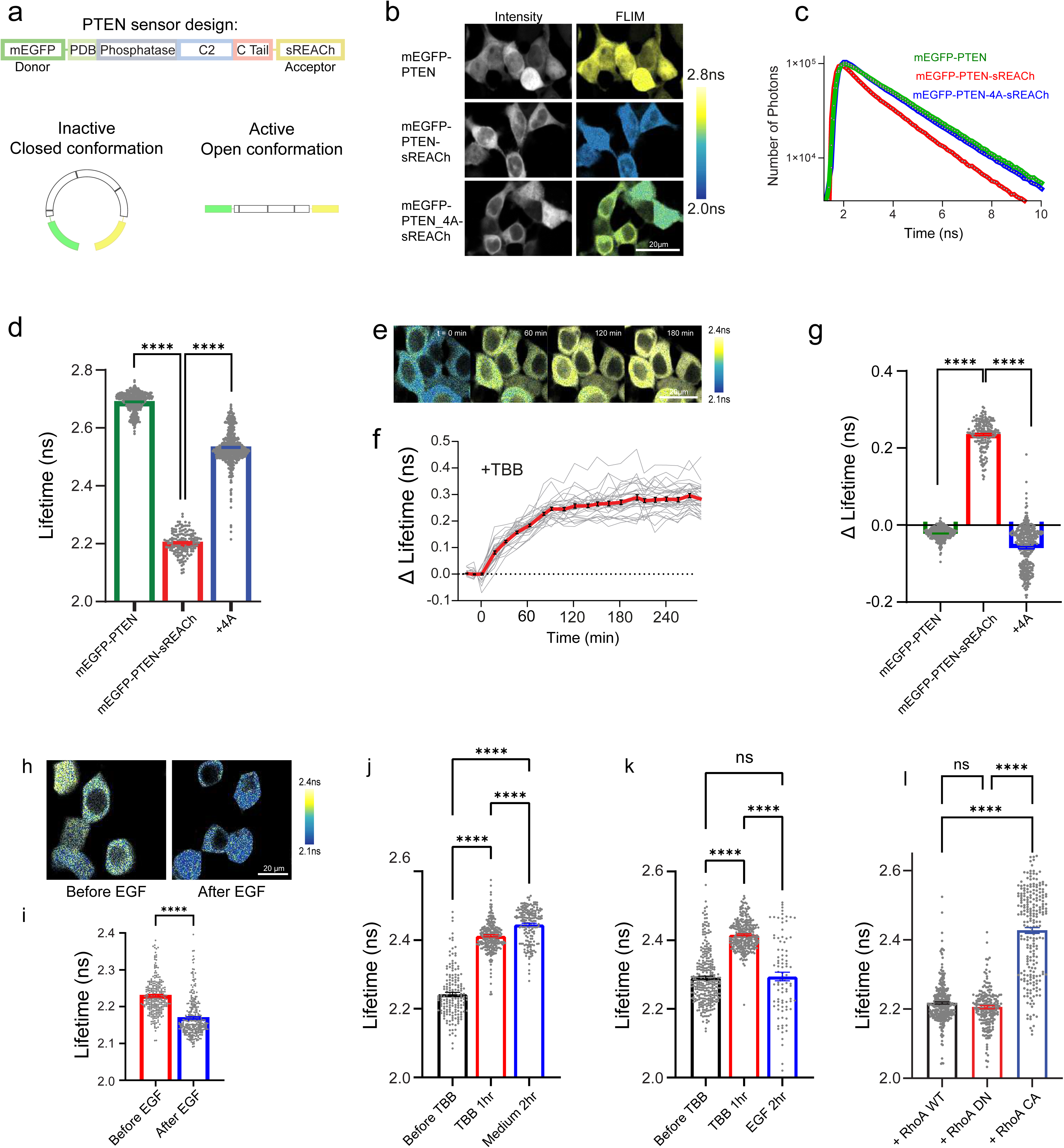
– Development of a new PTEN FRET/FLIM biosensor. (a) Schematic design of the FRET/FLIM-based mEGFP-PTEN-sREACh sensor. (b) Representative images of fluorescence intensity and pseudo coloured FLIM in HEK293 cells expressing mEGFP-PTEN (donor only), mEGFP-PTEN-sREACh or mEGFP-PTEN-4A-sREACh (4A mutant). Scale bar; 20 μm. (c) Representative fluorescence lifetime curves of HEK cells expressing mEGFP-PTEN, mEGFP-PTEN-sREACh or mEGFP-PTEN_4A-sREACh fitted with a double exponential decay. (d) Quantification of mean fluorescence lifetime in HEK cells expressing mEGFP-PTEN (2.69± 0.001 ns, n = 455), mEGFP-PTEN-sREACh (2.20±0.003 ns, n = 158) or mEGFP-PTEN_4A-sREACh (2.53±0.002 ns, n = 613). (e) Representative pseudo-coloured FLIM images of HEK cells expressing mEGFP-PTEN-sREACh at different time points following TBB application (50 μM), Scale bar; 20 μm. (f) Plot of changes in fluorescence lifetime over time of cells expressing mEGFP-PTEN-sREACh following TBB application. (g) Quantification of change in fluorescence lifetime following 3hr of TBB application of mEGFP-PTEN (−0.02±0.001 ns, n = 300), mEGFP-PTEN-sREACh (0.23±0.003 ns, n = 188) or mEGFP-PTEN_4A-sREACh (−0.06±0.003 ns, n = 372). (h) - (i) Pseudo-colored FLIM images and quantification of fluorescence lifetime of HEK293 transfected with mEGFP-PTEN-sREACh before (2.23±0.003 ns, n = 268) and after 100 ng/ml recombinant human EGF for 2hr (2.17±0.003 ns, n = 294). Scale bar; 20 μm. (j) Quantification of fluorescence lifetime of HEK293 transfected with mEGFP-PTEN-sREACh before TBB (2.24±0.005 ns, n = 166), after 1hr of TBB (2.41±0.003 ns, n = 277) and after cells were washed with medium (2.44±0.004 ns, n = 182). (k) Quantification of fluorescence lifetime of HEK293 transfected with mEGFP-PTEN-sREACh before TBB (2.29±0.004 ns, n = 290), after 1hr of 50 μM TBB (2.42±0.002 ns, n = 297) and after wash and application of 100ng/ml EGF for 2hr (2.29±0.012 ns, n = 90). ns p = 0.9233. (l) Quantification of fluorescence lifetime of HEK293 co-transfected with mEGFP-PTEN-sREACh and either WT RhoA (2.22±0.003 ns, n = 332), dominant negative (DN) RhoA (2.20± 0.004 ns, n = 201) or constituently active (CA) RhoA (2.43±0.008 ns, n = 225). ns p = 0.2565. Error bars represent SEM. Statistical difference was measured using one-way ANOVA followed by post-hoc Tukey’s multiple comparison test and unpaired two-tailed student t-test. **** p < 0.0001.

### PTEN sensor sensitivity to pathogenic mutation

Changes in PTEN signaling are associated with numerous pathologies. Previous work has demonstrated that distinct point mutations lead to PTEN loss of function ^25^. In the brain, PTEN mutations are one of the most frequently associated genetic causes of severe cases of autism spectrum disorders and macrocephaly ^22,23^. In addition, there is evidence that pathogenic mutations in PTEN lead to structural dysregulation ^45^ as well as reduced protein stability ^17^.

Accordingly, we set out to assess how different PTEN pathogenic mutations could be identified by our conformation-based PTEN biosensor in living cells. To this end, we introduced seven different point mutations, associated with forms of human cancer and autism, into the PTEN biosensor (**Fig. 2a**). We then expressed the different biosensor mutants in HEK293 cells and used 2pFLIM measurements to test for differences in fluorescence lifetime. Interestingly, we found that the point mutations exhibited a wide range of fluorescence lifetime (**Fig. 2b-c**). Some mutations caused a dramatic increase in lifetime, which indicates complete loss or de-stabilization of the compact closed form (mutations R173P, R130P, **Fig. 2b-c**).

**Figure 2.**
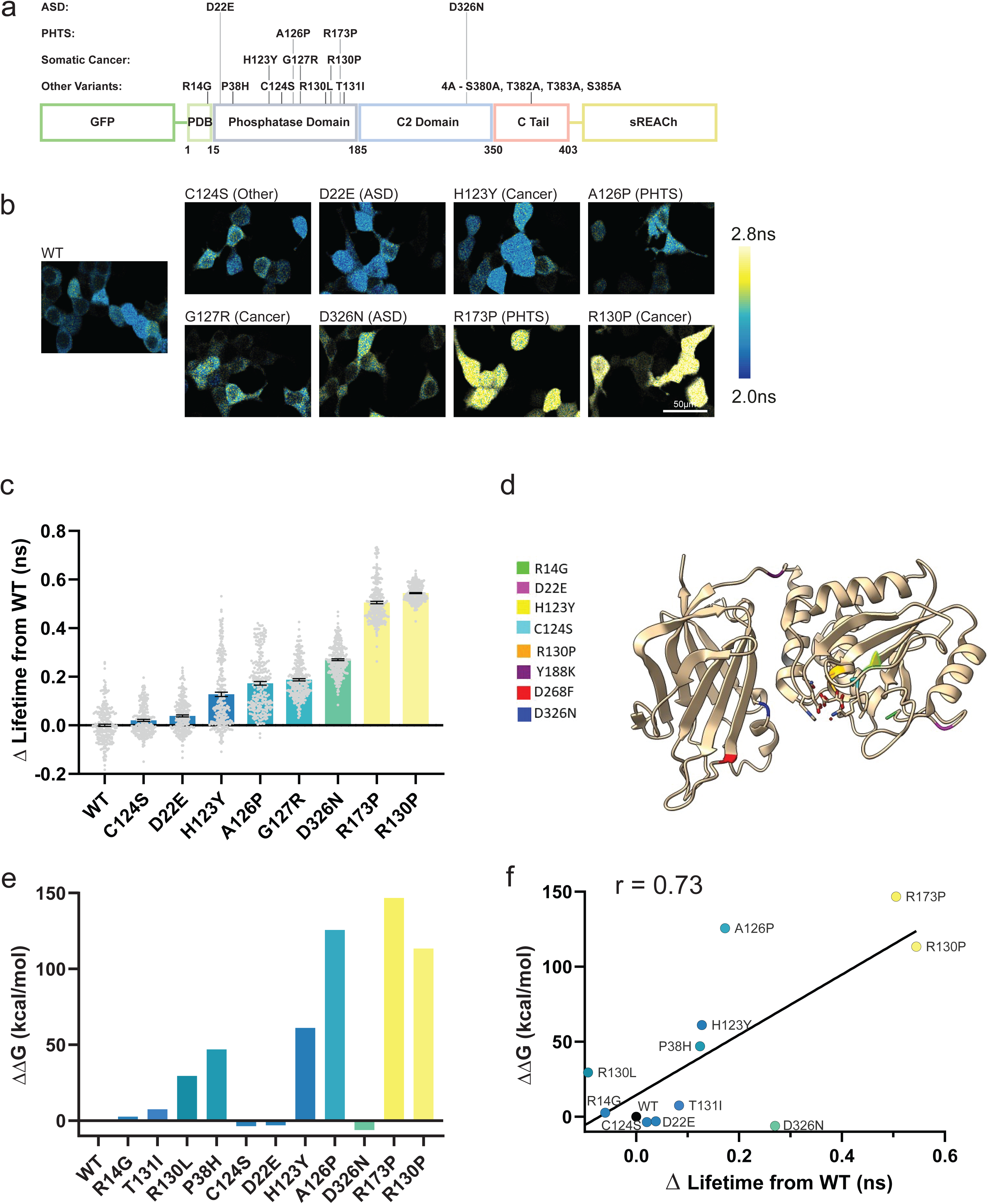
– PTEN sensor sensitivity to pathogenic mutation. (a) Schematic of PTEN mutants, their related pathology and their location along the protein, tested in the mEGFP-PTEN-sREACh biosensor. (b) Pseudo-colored FLIM images of HEK293 transfected with the mEGFP-PTEN-sREACh mutants. Scale bar; 50 μm. (c) Quantification of change in fluorescence lifetime of mEGFP-PTEN-sREACh containing mutations compared to WT version (0±0.005 ns, n = 258), C124S (p = 0.16, 0.02±0.004 ns, n = 225), D22E (****, 0.04±0.004 ns, n = 265), H123Y (****, 0.13±0.009 ns, n = 253), A126P (****, 0.18±0.007 ns, n = 250), G127R (****, 0.18±0.004 ns, n = 265), D326N (****, 0.27±0.004 ns, n = 259), R173P (****, 0.50±0.005 ns, n = 265) or R130P (****, 0.54±0.002 ns, n = 254). (d) Computer generated 3D model of PTEN based on its crystal structure, and the tested mutations’ locations along the protein. (e) Predicted ΔΔG of PTEN mutants compared to WT. R14G (2.78), T131I (7.52), R130L (29.59), P38H (47.05), C124S (−3.59), D22E (−2.99), H123Y (61.15), A126P (125.72), D326N (−6.03), R173P (146.70), R130P (113.44). Numbers represent units of kcal/mol. (f) Correlation between ΔΔG of PTEN mutants and their corresponding measured changes in fluorescence lifetime of mEGFP-PTEN-sREACh compared to WT (p = 0.007, r = 0.73). Error bars represent SEM; statistical difference for (c) was measured using one-way ANOVA followed by post-hoc Tukey’s multiple comparison test. Correlation was measured using two-tailed Pearson test and plotted using simple linear regression. One computer predicted outlier (G127R) was found and removed using Grubbs test for outliers (Alpha = 0.01). **** p < 0.0001.

As an orthogonal approach, we evaluated the effect of each mutation on protein stability by calculating the free energy difference between the wild-type and the different mutants. For this purpose, we relied on the published crystal structure of PTEN (PDB 1D5R) ^46^ and used the residue scanning and mutations module in the Schrodinger BioLuminate suite. **Figure 2e** displays a bar graph with the predicted ΔΔG, i.e., the difference between the ΔG values. This parameter represents the energy difference between the folded and unfolded state of each protein. Positive values for ΔΔG indicate protein destabilization relative to the wild-type. As expected, most of the mutations exhibit reduced stability (**Extended fig. 2**). Interestingly, we found a strong correlation between the fluorescence lifetime differences and the predicted stability caused by individual point mutations (**Fig. 2f**, r = 0.73).

Altogether, our FRET-based approach detects structural abnormalities caused by pathogenic PTEN mutations with high sensitivity. This makes the PTEN biosensor suitable for the systematic characterization of new PTEN mutations.

### Optimization of the PTEN biosensor to minimize interference with endogenous signaling

Following our initial characterization of the PTEN biosensor, we set out to further utilize this approach *in vivo*. One possible limitation is potential interference with cellular function due to overexpression of PTEN. Indeed, previous studies reported that PTEN overexpression leads to structural loss of dendritic spines, as well as morphological and physiological perturbations ^14,17,31^. To overcome these potential caveats, we screened different PTEN LOF point mutations, focusing on four different mutants that displayed minimal biochemical activity but maintained good protein stability in a recent PTEN mutant screen ^17^. We expressed these different mutations in HEK293 cells and tested their basal lifetime and conformational dynamics following activation of PTEN induced by TBB (**Fig. 3a-c**). Despite a range of basal conformations and changes, we identified the R14G mutant as similar to the PTEN WT with respect to lifetime, expression, and brightness (**Extended fig. 3a).** This mutant also displayed a significant magnitude of TBB induced conformational change (**Fig. 3b-c**), and reversible changes in lifetime similar to those of the WT sensor (**Fig. 3d-g**). We decided to continue to characterize this variant, since we reasoned that a PTEN biosensor harboring the R14G mutation should retain conformational sensitivity as a proxy for activity but display minimal downstream signaling due to loss of catalytic activity ^17^. We therefore tested how overexpression of the PTEN biosensor affects cellular morphology as assessed by cytosolic co-expression of a fluorescent cell-fill (CyRFP). The results indicated that TBB induction of PTEN activity reduced cell-size in control cells (**Extended fig. 3b-c)**. Interestingly, co-expression of the WT PTEN sensor significantly decreased HEK293 cell size, with no further reduction after application of TBB. However, overexpressing the R14G mutant biosensor did not cause any changes in cell size, and the reduction in size was similar to that of control cells after TBB application (**Extended fig. 3b-c)**. Previous work has demonstrated that PTEN KO leads to a dramatic increase in neuronal soma size and increases dendritic spine density ^9,47^. To determine the impact of overexpressed PTEN R14G biosensor on neuronal morphology in vivo in the intact brain, we performed in utero electroporation (IUE) ^48^ of plasmids containing SpCas9 and a gRNA^49^ targeting PTEN exon 6, together with CyRFP (**Fig. 3h**). This approach was previously shown to knock-out PTEN ^50^. We validated that the gRNA indeed effectively reduced mouse PTEN (**Extended fig. 3d-e**). Following IUE, adult mice underwent cranial window surgery followed by in vivo imaging after postnatal day 30. The results indicate that L2/3 neurons expressing spCas9/gRNA exhibit morphological features of hypertrophic soma, in contrast to control cells that were electroporated with a scrambled gRNA sequence (**Fig. 3i-j**). To characterize the PTEN R14G biosensor, we tested the structural properties of L2/3 neurons expressing PTEN WT or the R14G sensor on the background of spCas9/gRNA for PTEN. The results reveal that overexpression of WT PTEN completely restored the soma size to control levels **(Fig. 3i-j**). However, overexpression of the mutant R14G sensor could not restore soma size. Characterization of the synaptic density of apical L2/3 dendrites in the same experiments indicated that PTEN gRNA leads to an increase in spine density compared to a scrambled control. Overexpression of WT PTEN restored spine density to control levels, while overexpression of the mutant R14G containing biosensor did not change the spine density compared to KO **(Fig. 3k-l)**.

**Figure 3.**
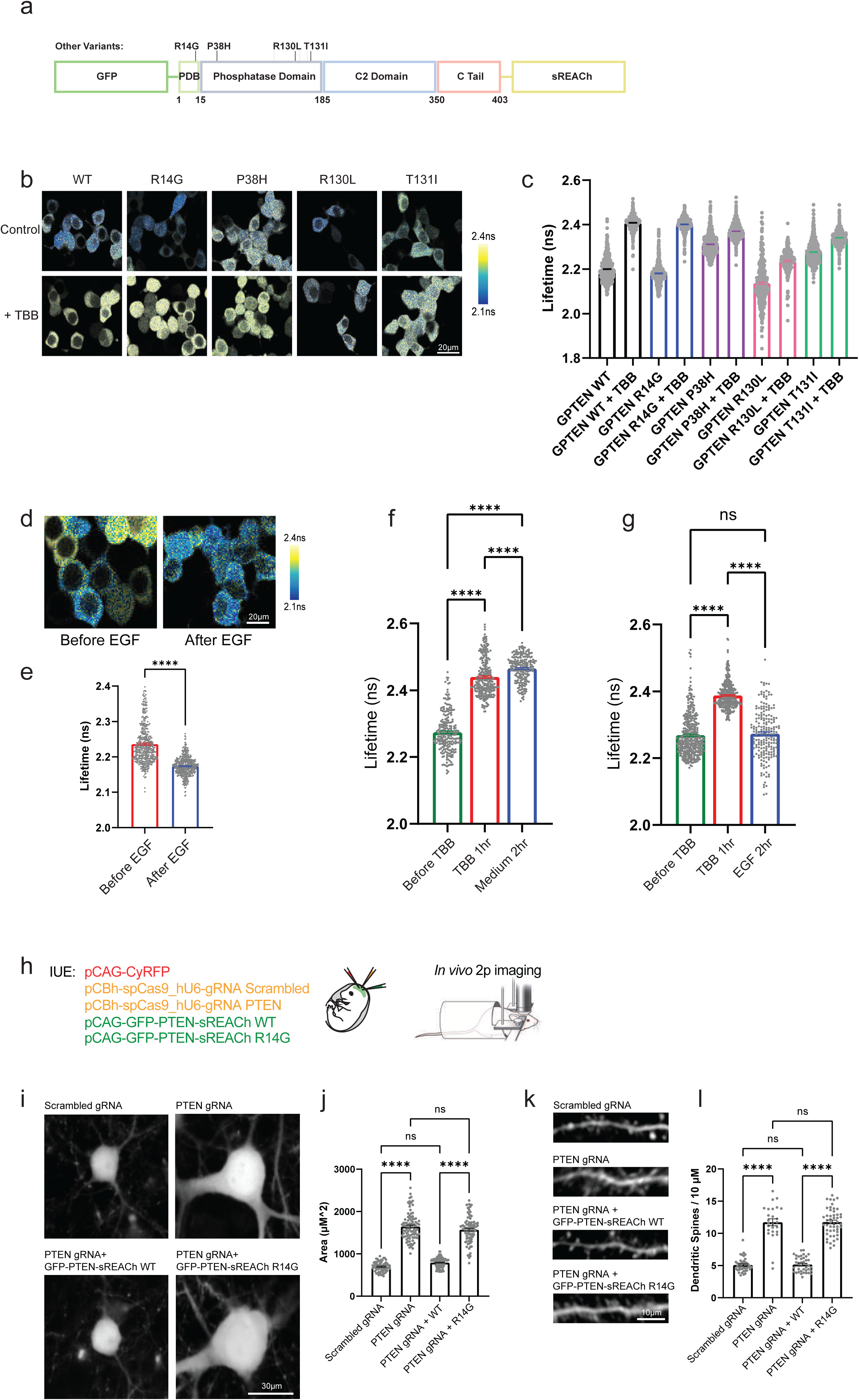
– Optimization of PTEN biosensor to minimize interference with endogenous signaling. (a) Schematic of the PTEN biosensor mutant candidates. (b) Pseudo-colored FLIM images of HEK293 transfected with the mEGFP-PTEN-sREACh mutants. Scale bar; 20 μm. (c) Quantification of basal fluorescence lifetime of the different mEGFP-PTEN-sREACh candidate mutants, and following 3hr of TBB (50 μM) application, WT (****, 2.20±0.002 ns, n = 502, and 2.41±0.002 ns, n = 295), R14G (****, 2.18±0.002 ns, n = 478, and 2.40± 0.002 ns, n = 399) P38H (****, 2.31±0.002 ns, n = 480, and 2.37±0.002 ns, n = 499), R130L (****, 2.14±0.005 ns, n = 352, and 2.24±0.004 ns, n = 202), T131I (****, 2.28±0.002 ns, n = 495, and 2.34±0.003 ns, n = 503) for control and after TBB application, respectively. (d) - (e) Pseudo-colored FLIM images and quantification of fluorescence lifetime of HEK293 transfected with mEGFP-PTEN-sREACh containing the R14G mutation (G-PTEN) before (2.23±0.003 ns, n = 442) and after 100 ng/ml recombinant human EGF (2.17n ± 0.001 ns, n = 420). Scale bar; 20 μm. (f) Quantification of fluorescence lifetime of HEK293 transfected with G-PTEN before (2.27± 0.004 ns, n = 232), after 1hr of 50 μM TBB application (2.44±0.003 ns, n = 276), and following wash in medium for 2hr (2.46±0.003 ns, n = 210). (g) Quantification of fluorescence lifetime of HEK293 transfected with G-PTEN before (2.27± 0.003 ns, n = 374), after 1hr of 50 μM TBB (2.39±0.002 ns, n = 357) and following wash in 100ng/ml recombinant human EGF application for 2hr (2.27±0.005 ns, n = 194). ns p = 0.5980. (h) Schematic of In-utero electrophoresis (IUE) injections in mice brains and in vivo 2pFLIM of the adult mice. (i) Representative 2p in vivo images of CyRFP labeled L2/3 cells, co-expressing spCas9 and scrambled control gRNA (Scr), or a PTEN targeted gRNA. In addition, rescue experiments representative images for cells co-expressing mEGFP-PTEN-sREACh WT or R14G. Scale bar; 30 μm. (j) Quantification of neuron soma area in μm^2^ indicated by CyRFP, for neurons co-expressing scrambled guide RNA (696.6±14.72, n = 62), PTEN gRNA (1646±32.77, n = 93), PTEN gRNA+mEGFP-PTEN-sREACh WT (791.5±14.5, n = 78) and PTEN gRNA + mEGFP-PTEN-sREACh R14G (1634±28.87, n = 74). ns p = 0.0787 and p = 0.1554 for Scrambled gRNA compared to PTEN gRNA + WT and for PTEN gRNA compared to PTEN gRNA + R14G respectively. (k) Representative 2p in vivo images of fluorescently labeled apical denrites expressing CyRFP, and co-expressing spCas9 and scrambled guide RNA or PTEN targted gRNA, rescued with either mEGFP-PTEN-sREACh WT or R14G. Scale bar; 10 μm. (l) Quantification of spine density per 10 μm for dendrites expressing CyRFP and scrambled guide RNA (5.087±0.161, n = 43), PTEN gRNA (11.72±0.53, n = 26), PTEN gRNA + mEGFP-PTEN-sREACh WT (5.18±0.18, n = 40), and PTEN gRNA + mEGFP-PTEN-sREACh R14G (11.77±0.29, n = 49). ns p = 0.9956 and p = 0.9995 for Scrambled gRNA compared to PTEN gRNA + WT and for PTEN gRNA compared to PTEN gRNA + R14G respectively. Error bars represent s.e.m; statistical differences for (c, f, g, i, k) were measured using one-way ANOVA followed by post-hoc Tukey’s multiple comparison test. Statistical difference for (e) was measured using unpaired two-tailed student t-test. **** p < 0.0001.

In conclusion, our results validate the R14G variant (termed as G-PTEN) as a robust probe for the assessment of PTEN activity state, but with a minimal impact on endogenous PTEN signaling.

### Development and utilization of a transgenic strain of *C. elegans* designed to detect PTEN signaling

Our next step involved the development of a small experimental model to assess the G-PTEN sensor in an intact nervous system. Given the wide conservation of PTEN across species, we selected *C. elegans*, where the PTEN homolog *daf-18* plays a key role in metabolic, developmental, and longevity pathways ^51–53^. We therefore cloned the PTEN R14G sensor under the neuronal worm promoter P*snb-1* ^54^ and injected it into *C. elegans* to create a transgenic worm with pan-neuronal expression **(Fig. 4a)**. We then conducted 2pFLIM measurements on whole anesthetized worms. Since CK2 is conserved in the worm ^55^, we validated the functionality of the PTEN sensor using TBB as an effective pharmacological agent to induce PTEN activation. Adding TBB to the worms’ food, led to a robust increase in neuronal PTEN lifetime, which corresponds to increased PTEN activity (**Fig. 4b-c**). In *C. elegans*, *daf-2*, the Insulin-like Growth Factor 1 (IGF1) homologue, serves a critical role in longevity and proteostasis ^56,57^. This signaling pathway is well conserved in mammals where both IGF1 and PTEN are vital regulators of longevity, proteostasis, and cell-metabolism ^33,58^. However, the ability of *daf-2* to regulate PTEN activity directly throughout development has never been investigated. To resolve this issue, we used 2pFLIM and the PTEN biosensor to characterize the PTEN activity throughout the stages of worm larval development. The results indicate that PTEN undergoes a gradual increase in activity across the L1 to adult worm stages, as assessed by an increase in the basal lifetime (**Fig. 4d-e**). The increase in activity reached a plateau around L4, suggesting an association with neuronal maturity at the L4 stage. Interestingly, inhibition of *daf-2* by RNAi ^59^, revealed that *daf-2* downregulation induces a significant increase in PTEN activity, which is maintained throughout development (**Fig. 4d, f-g)**. Overall, these results validate the utility of the PTEN biosensor for direct visualization of PTEN signaling in *C. elegans*, and suggest a direct interplay between *daf-2* and PTEN, which have both been previously implicated as critical regulators of longevity ^33,52^.

**Figure 4.**
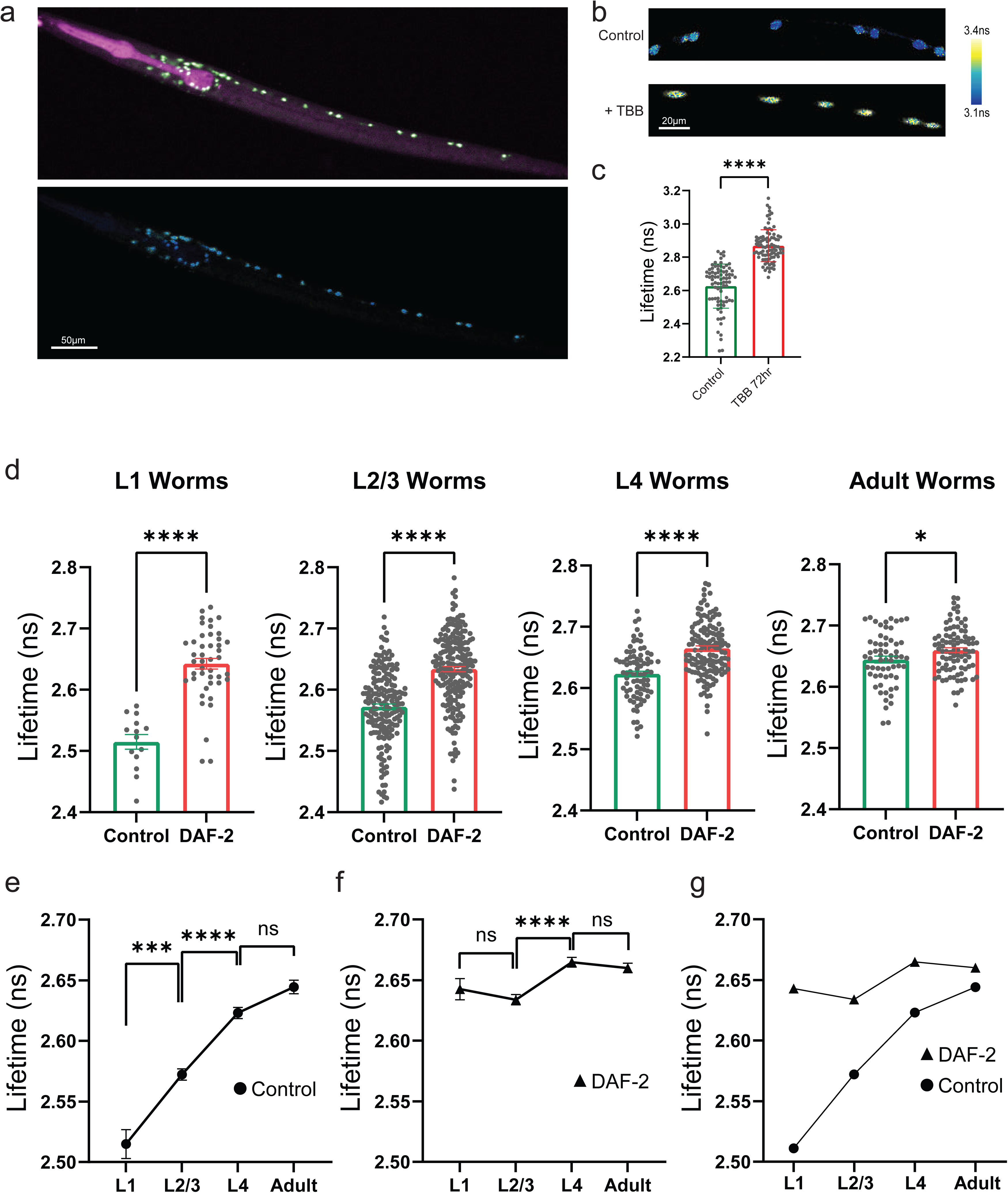
– Development and utilization of a transgenic *C. elegans* to detect PTEN signaling. (a) Top-Representative fluorescent image of labeled *C. elegans* with pan-neuronal G-PTEN (green) and pharyngeal marker (magenta). In the bottom-pseudo-colored FLIM image of the same field of view. Scale bar; 50 μm. (b) – (c) Pseudo-colored FLIM images and quantification in neurons of *C. elegans* expressing pan-neuronal G-PTEN before (2.63±0.02 ns, n = 81) and after 72hr of 500uM TBB (2.87±0.01 ns, n = 90). Scale bar; 20 μm. (d) Quantification of fluorescence lifetime of G-PTEN at each stage of larval development, with or without *daf-2* RNAi. Larvae were measured at L1 stage (2.51±0.014 ns, n = 14, and 2.64±0.01 ns, n = 44), L2/3 stage (2.57±0.005 ns, n = 184, and 2.63±0.004 ns, n = 204), at L4 stage (2.62±0.005 ns, n = 82, and 2.66±0.004 ns, n = 141) or at adulthood (p = 0.02, 2.64±0.006 ns, n = 61, and 2.66±0.004 ns, n = 90) without or with *daf-2* RNAi, respectively. (e) – (g) Quantification of fluorescence lifetime of G-PTEN transgenic *C. elegans* neurons comparing each stage of larval development, for control group or with *daf-2* RNAi. In (e) *** p = 0.0005 and ns p = 0.1004. In (f) ns p = 0.7504 and p = 0.9099 for L1 compared to L2/3 and for L4 compared to Adult, respectively. Error bars represent s.e.m; statistical differences for (d, e) were measured using unpaired two-tailed student t-test. Statistical differences for (f, g) were measured using one-way ANOVA followed by post-hoc Tukey’s multiple comparison test. **** p < 0.0001.

### In vivo imaging of PTEN in the mouse brain

In order to measure PTEN activity in the living mouse brain, we used IUE to sparsely express the G-PTEN sensor in the mouse L2/3 motor and somatosensory cortex ^48^. We performed cranial window surgery ^60^ in adult electroporated mice following postnatal day 45 **(Fig. 5a**). We found that the combination of sparse expression of G-PTEN and in vivo 2pFLIM allowed monitoring of PTEN activity at superb subcellular resolution in the neuronal cell body, individual dendrites, and in dendritic spines **(Fig. 5b)**. Overall, the results indicated that the G-PTEN lifetime is elevated in the cell body compared to dendritic regions, which suggests compartmentalized PTEN activity in vivo **(Fig. 5c)**. We then tested whether genetic manipulation of key genes in the PTEN signaling complex could lead to changes in PTEN activity in vivo. Both Insulin-like Growth Factor 1 Receptor (IGF1R) and Tuberous Sclerosis complex subunit 2 (Tsc2) are conserved genes that are vital for neuronal function, and act upstream and downstream of PTEN in its neuronal signaling complex ^8^. To test the effect of their perturbation on PTEN, we co-expressed G-PTEN with a plasmid expressing spCas9 and gRNA, targeted towards IGF1R, and Tsc2. We validated the efficiency of CRISPR/Cas9 based targeting by perturbing IGF1R and Tsc2 and then testing for the induction of phosphorylated S6 (pS6), a common downstream target in the signaling pathway ^9^ **(Extended fig. 4a-b, e-f)**. We also validated that the IGF1R gRNA indeed effectively reduced IGF1R expression (**Extended fig. 4c-d**). Using in vivo 2pFLIM of G-PTEN, we found that PTEN activity increases in the soma following IGF1R and Tsc2 gRNA, as evidenced by a longer lifetime, compared to G-PTEN with a scrambled gRNA (**Fig. 5d-e**). However, morphological changes were characterized by an increase in cell body size following Tsc2 KO, and a decrease following IGF1R (**Fig, 5d, f**). Analysis of changes in the dendritic PTEN lifetime revealed distinct changes in dendritic PTEN activity: PTEN activity in the Tsc2 KO dendritic shaft was unchanged, whereas IGF1R KO led to an increase in PTEN activity. In contrast, IGF1R KO did not alter the synaptic PTEN lifetime, which was significantly decreased by Tsc2 KO (**Fig. 5g-h**). These changes in PTEN activity were accompanied by structural changes: dendritic spine density was increased compared to controls after Tsc2 KO, while IGF1R KO led a decrease in spine density (**Fig. 5i**). Importantly, in vivo imaging of PTEN activity allowed us to identify differences in subcellular PTEN activity in the synaptic and somatic compartments following manipulation of upstream or downstream signaling. These changes are associated with distinct structural changes and confirm the critical role of PTEN activity and the close association with neuronal and synaptic trophism.

**Figure 5.**
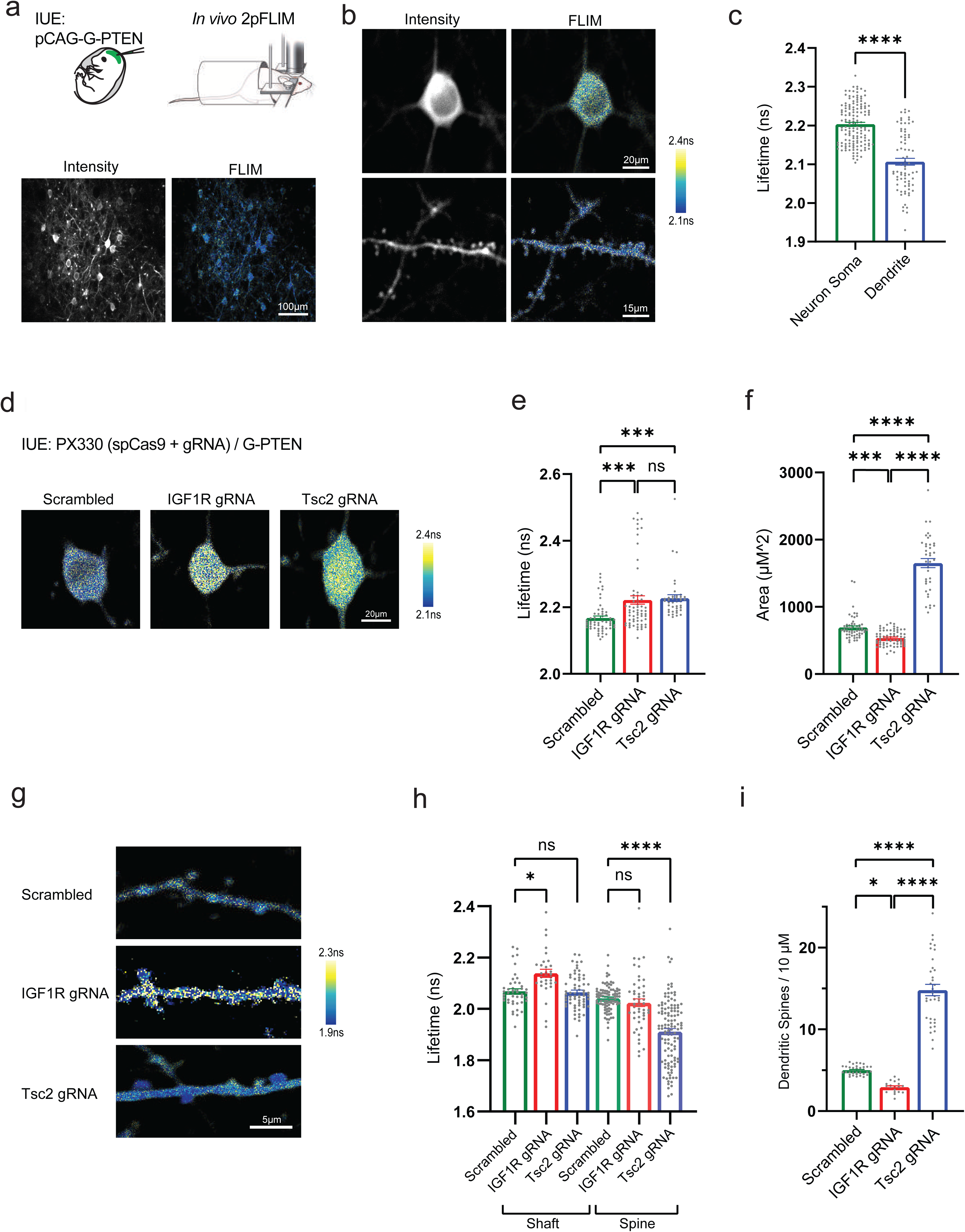
– In vivo imaging of PTEN in the mouse brain. (a) Top-schematic of In-utero electroporation (IUE) followed by in vivo 2pFLIM in the adult mouse brain. Bottom-Representative widefield images of fluorescent intensity and FLIM of L2/3 cells expressing G-PTEN. Scale bar; 100 μm (b) Representative high magnification images of fluorescence intensity and pseudo-colored lifetime of G-PTEN expression in L2/3 soma (top) and dendrite (bottom). Scale bars; Soma - 20 μm, Dendrite - 15 μm. (c) Quantification of fluorescence lifetime in G-PTEN expressing neuronal somas (2.20±0.004 ns, n = 142) and dendrites (2.11±0.009 ns, n = 75) in 4 mice. (d) Representative 2p in vivo images of L2/3 neuronal somas co-expressing G-PTEN, spCas9 and either scrambled control gRNA (Scr), IGF1R gRNA or a Tsc2 targeted gRNA. Scale bar; 20 μm. (e) Quantification of G-PTEN fluorescence lifetime in L2/3 neuronal somas targeted with either scrambled control gRNA (2.17±0.006 ns, n = 56), an IGF1R targeted gRNA (p = 0.0005, 2.22±0.013ns, n = 64) or a Tsc2 targeted gRNA (p = 0.0006, 2.22±0.01ns, n = 40). ns p = 0.9263. (f) Quantification of L2/3 soma area in μm^2^ of neurons expressing scrambled gRNA (693.1±22.86, n = 57), IGF1R gRNA (p = 0.0009, 537.5±12.87, n = 70) or Tsc2 gRNA (1652±66.87, n = 39). (g) Representative 2p in vivo images of L2/3 neuronal dendrites co-expressing G-PTEN, spCas9 and scrambled control gRNA (Scr), IGF1R gRNA or a Tsc2 gRNA. Scale bar; 5 μm. (h) Quantification of G-PTEN fluorescence lifetime in L2/3 neurons targeted with either scrambled control gRNA (2.07±0.009 ns, n = 45, and 2.04±0.005 ns, n = 94), IGF1R gRNA (2.14±0.016 ns, n = 31, and 2.02±0.015 ns, n = 49) or Tsc2 gRNA (2.07±0.008ns, n = 65, and 1.91±0.012ns, n = 120) for their dendritic shafts and spines, respectively. * p = 0.0188. ns p > 0.9999 and p = 0.8882 for Scrambled Shaft compared to Tsc2 gRNA Shaft and for Scrambled Spine compared to IGF1R gRNA Spine, respectively. (i) Quantification of spine density per 10 μm for G-PTEN neurons expressing scrambled control gRNA (5±0.09, n = 35), IGF1R targeted gRNA (p = 0.0197, 2.93±12.87, n = 19) or a Tsc2 targeted gRNA (14.8±0.7, n = 35). Error bars represent s.e.m; statistical difference for (c) was measured using unpaired two-tailed student t-test. Statistical differences for (e, f, h, i) were measured using one-way ANOVA followed by post-hoc Tukey’s multiple comparison test. **** p < 0.0001.

### Dual imaging of PTEN activity using a red-shifted PTEN sensor

Next, we set out to develop a red-shifted variant of the PTEN sensor. Our strategy was to utilize a previously developed FRET pair optimized for 2pFLIM, namely mCyRFP2 and mMaroon, as donor and acceptor ^37,42^. We therefore replaced mEGFP and sREACh in the G-PTEN sensor with mCyRFP2 and mMaroon1, respectively (**Fig. 6a**). This new variant, named R-PTEN, retains sensitivity and specificity, as the 4A mutation increases the basal lifetime (**Fig. 6b-c**). TBB application led to a significant increase in lifetime (**Fig. 6d**), and EGF application reduced the lifetime **(Fig. 6e**). Following initial characterization, we tested the application of in vivo imaging of R-PTEN. For this purpose, we prepared adeno associated Viruses (AAV) of R-PTEN with a neuronal Synapsin promoter that limits expression to excitatory cells. We utilized a recently established approach for codon diversification ^61^ of the acceptor mMaroon to avoid AAV based recombination of FRET sensors. The unique properties of mCyRFP2 allowed us to co-express and simultaneously image it alongside GFP based GCaMP8s ^62^ (**Fig. 6f-g**). This is due to the long-stoke shift of mCyRFP, which allows separation of the emission spectra after single wavelength excitation of both GFP and R-PTEN. We reasoned that this combination would allow us to monitor ongoing neuronal activity levels and PTEN activity at the same time in individual neurons in vivo. In vivo imaging of R-PTEN and GCaMP8s in the L2/3 somatosensory cortex, revealed calcium transients of GCaMP8s in the green channel (**Fig. 6g-h, Supplementary movie 2**) while monitoring the R-PTEN lifetime, without green to red channel crosstalk. Spontaneous neuronal activity, which correlates with functional connectivity, was measured in order to validate and examine the properties of R-PTEN expressing cells compared to nearby non-expressing cells. Notably, there was no significant difference in ongoing spontaneous activity between R-PTEN positive and negative cells in the same mice (**Fig 6i**). These results further confirm the validity of our general approach using the R14G mutation to decrease downstream signaling activity, which preserves normal physiological properties even after long-term in vivo expression of the sensor. We tested the link between PTEN activity levels and spontaneous activity in the same population of L2/3 cells in vivo. We found that overall spontaneous activity is inversely correlated with the PTEN lifetime across cortical cells (r = −0.21, **Fig. 6j**). Our overall results confirm the usefulness of the PTEN sensor for in vivo imaging of PTEN activity and the ability to use it in a combination of different biosensors to probe neuronal signaling and activity in vivo.

**Figure 6.**
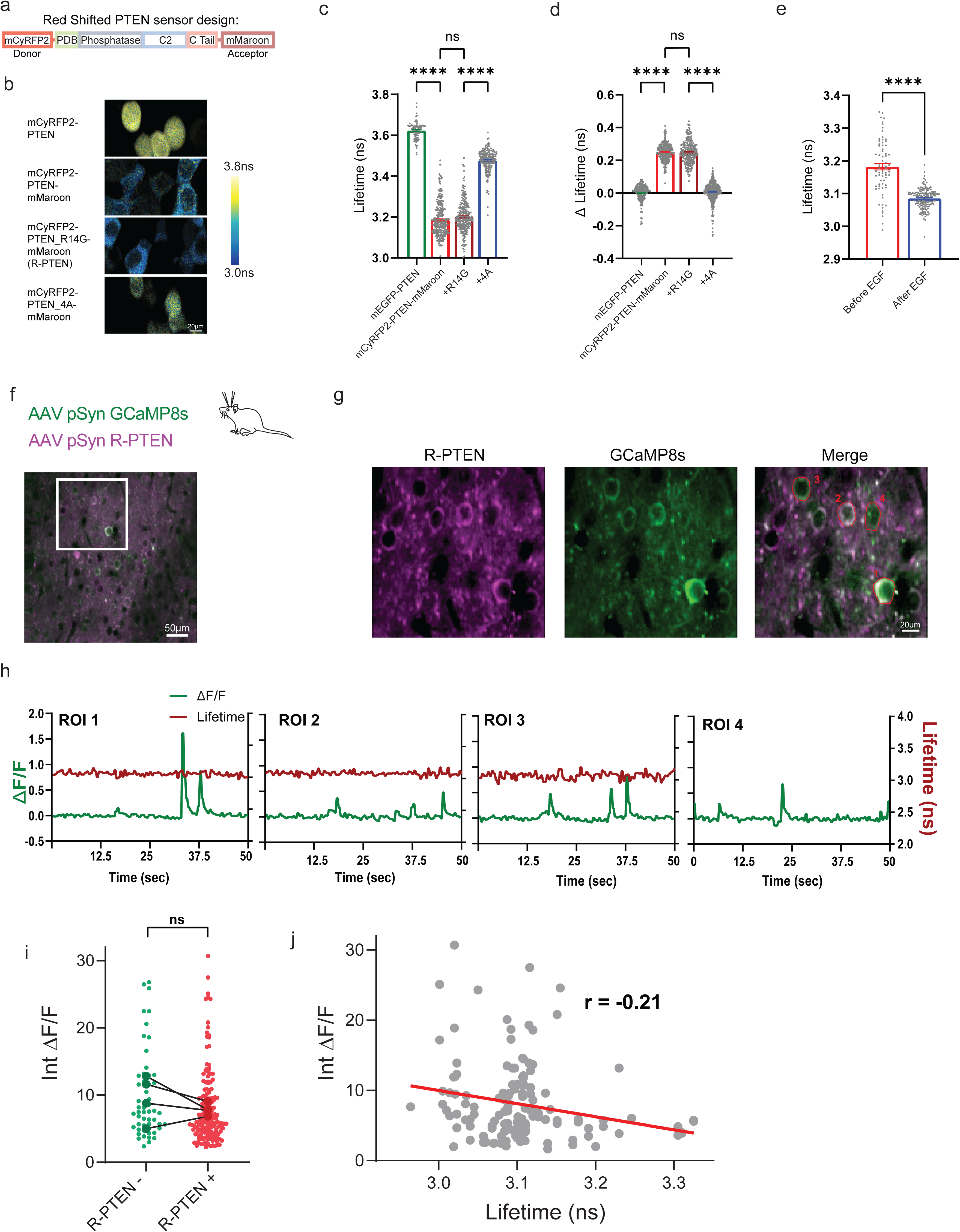
– Dual imaging of PTEN activity using a red-shifted PTEN sensor. (a) Schematic design of the FRET/FLIM-based mCyRFP2-PTEN-mMaroon sensor. (b) Representative images of pseudo colored FLIM in HEK293 cells expressing mCyRFP2-PTEN (donor only), mCyRFP2-PTEN-mMaroon, mCyRFP2-PTEN-mMaroon R14G (R-PTEN) or mCyRFP2-PTEN-4A-mMaroon (4A mutant). Scale bar; 20 μm. (c) Quantification of mean fluorescence lifetime in HEK cells expressing mCyRFP2-PTEN (3.62±0.004 ns, n = 88), mCyRFP2-PTEN-mMaroon (3.19±0.005 ns, n = 279), mCyRFP2-PTEN-mMaroon R14G (3.20±0.005 ns, n = 220) or mCyRFP2-PTEN-4A-mMaroon (3.48±0.004 ns, n = 199). ns p = 0.0941. (d) Quantification of change in fluorescence lifetime following 3hr of TBB application in mCyRFP2-PTEN (0.01±0.004 ns, n = 200), mCyRFP2-PTEN-mMaroon (0.25±0.003 ns, n = 292), mCyRFP2-PTEN-mMaroon R14G (0.25±0.004 ns, n = 234) or mCyRFP2-PTEN-4A-mMaroon (0.01±0.003 ns, n = 400). ns p = 0.9987. (e) Quantification of fluorescence lifetime of HEK293 transfected with R-PTEN before (3.18±0.009 ns, n = 74) and after 100 ng/ml recombinant human EGF for 2hr (3.085±0.002 ns, n = 155). (f) Schematic of AAV injections in p3-p4 mice and a low magnification representative in vivo image of R-PTEN (magenta) and GCaMP8s (green) in L2/3 somatosensory cortex in a p28 mouse. Scale bar; 50 μm (g) In vivo high magnification images of the boxed region in (f) of individual L2/3 cells expressing R-PTEN (magenta), GCaMP8s (green) and their merged image. Scale bar; 20 μm. (h) Time course of changes in ΔF/F (GCaMP8s, green) and lifetime in the red channel (R-PTEN, red) of cells marked in (g) in co-expressing cells (ROI1-3), and in a cell expressing GCaMP8s only (ROI4). (i) Comparison of summed integrated ΔF/F (arbitrary units) in R-PTEN negetive neurons (n = 53) and R-PTEN positive neurons (n = 164), paired for each mouse (n = 4 mice, p = 0.32). (j) Correlation between fluorescence lifetime and cumulative summed integrated ΔF/F in R-PTEN+ neurons (p = 0.008, r = −0.21). Error bars represent SEM; Statistical differences for (c, d) were measured using one-way ANOVA followed by post-hoc Tukey’s multiple comparison test. Statistical difference for (e) was measured using unpaired two-tailed student t-test. Statistical difference for (i) was measured using paired two-tailed student t-test. Correlation was measured using two-tailed Pearson test and graphically plotted using simple linear regression. **** p < 0.0001.

### Simultaneous cell-type specific in vivo imaging of PTEN signaling

Lastly, we examined the use of the PTEN sensor to explore cell specific PTEN activity in the intact brain. Previous studies have reported differences in PTEN signaling in inhibitory cells ^18^, hippocampal dentate gyrus cells ^27^, and astrocytes ^10^. However, the precise contribution of PTEN signaling was inferred by cell-type specific PTEN KO. Since PTEN KO leads to structural and functional dysregulation, it is possible that PTEN confers a unified signaling tone across cells, but that this produces different phenotypical manifestations due to cell specific functions.

It is therefore critical to examine whether physiological PTEN activity is differentially regulated in distinct cell-types in the brain. To address this question using the G-PTEN biosensor, we first explored an experimental approach to monitor PTEN activity in L2/3 excitatory cells and cortical astrocytes using G-PTEN. The piggyBac vector allows genomic integration of a DNA transposon-based insertion into progenitor cells of the developing cortex ^63^. This can be used to label both L2/3 neurons and cortical astrocytes, since they are derived from the same common progenitors ^64^. IUE of G-PTEN under the control of a piggyBac transposon vector, together with a hyperactive piggyBac transposase (**Extended Fig. 5)** efficiently labeled L2/3 neurons and astrocytes with G-PTEN in the same mouse. Cortical neurons and astrocytes could be distinguished by their distinct morphology and anatomical position. Interestingly, while the basal PTEN lifetime was relatively consistent across L2/3 soma, astrocytes displayed heterogenous values, which were on average lower than neuronal levels (**Extended Fig. 5b)**. Differences in basal PTEN activity could be attributed to the complex morphological and proliferation rate of astrocytes.

Then, we set out to determine PTEN activity in excitatory and inhibitory cells in the somatosensory cortex. Previous studies have illustrated important roles for PTEN signaling in both excitatory and inhibitory cells, but it remains unclear whether PTEN activity is differentially regulated under physiological conditions. In the cortex, excitatory and inhibitory cells are intermingled and therefore we chose to use a dual-color genetic strategy involving R-PTEN and G-PTEN. First, we constructed an AAV with a double FLOXed Cre recombinase dependent expression of G-PTEN (**Extended fig. 6**). We used an AAV with a human Synapsin promoter expressing R-PTEN to label excitatory cells (**Extended fig. 6c**). We then used a parvalbumin (PV) Cre line, and co-injected AAVs for R-PTEN and G-PTEN. This combination allows us to target excitatory cells and PV inhibitory cells in the same field of view simultaneously (**Fig. 7a-b**). Using this approach, we set out to examine whether sensory experience alters cell-type specific PTEN activity in the brain. Following AAV injections in neonatal mice, we performed whisker deprivation of the contralateral whisker for 2 weeks during the critical period and into adulthood (p11-p28) ^65^. Previous studies have reported that this manipulation profoundly alters the excitatory to inhibitory ratio (E/I) in the somatosensory cortex ^66,67^. Comparing the lifetime of G-PTEN in PV cells and R-PTEN in excitatory cells in control mice with intact whiskers, and mice after contralateral whisker trimming **(Fig. 7c-d)**, revealed cell-specific changes in PTEN activity. Excitatory PTEN activity was slightly reduced following whisker deprivation. Conversely, inhibitory PTEN activity increased following deprivation (**Fig. 7e**). We can therefore conclude that sensory deprivation leads to a distinct reduction of E/I levels of PTEN signaling (**Fig. 7f)**.

**Figure 7.**
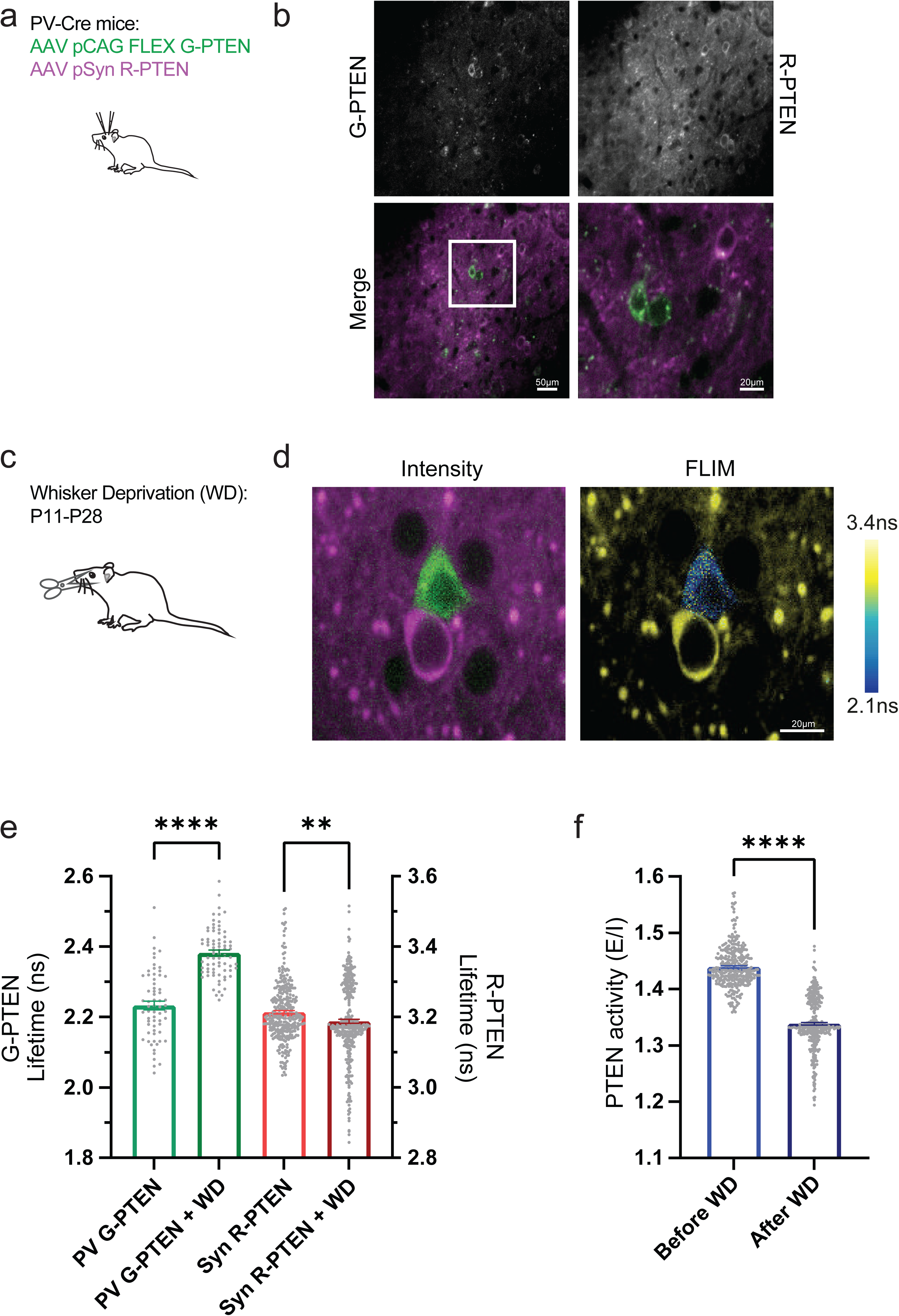
– Simultaneous in vivo imaging of PTEN signaling in excitatory and inhibitory cells. (a) Schematic of cell-type specific AAV injections in neonatal mice. (b) Representative images of PV cells labeled with Cre dependent G-PTEN (left / green) and excitatory cells labeled with R-PTEN (left / magenta) in the somatosensory cortex of a p28 mouse. Widefield scale bar; 50 μm. Zoomed scale bar; 20 μm. (c) Schematic of contralateral whisker deprivation (WD) between p11-p28. (d) Representative intensity and FLIM images of excitatory (magenta) and inhibitory (green) cells after WD. Scale bar; 20 μm. (e) Quantification of fluorescence lifetime before (2.23±0.012ns, n = 64, and 3.214±0.005ns, n = 317) and after WD (2.38±0.008ns, n = 76, and 3.19±0.006ns, n = 364) for Inhibitory G-PTEN expressing cells (left / green) and Excitatory R-PTEN expressing cells (right / red), respectively. ** p = 0.0026. (f) Excitatory/Inhibitory ratio (E/I) of R-PTEN/G-PTEN fluorescence lifetime before (1.44±0.002ns, n = 317) and after WD (1.34±0.002ns, n = 364). Error bars represent SEM; statistical difference for (e) was measured using one-way ANOVA followed by post-hoc Tukey’s multiple comparison test. Statistical difference for (f) was measured using unpaired two-tailed student t-test. **** p < 0.0001.

Altogether, simultaneous imaging of PTEN signaling in excitatory and inhibitory cells revealed cell-type specific dynamics following sensory experience. Future work could allow to identify further unique PTEN signaling dynamics within cell-specific neuronal cells, as well as astrocytes and microglia.

## Discussion

Here, we describe the development and validation of a novel biosensor that can dynamically monitor PTEN signaling in intact biological systems – cell lines, intact tissues, and whole organisms. The combination of a FRET biosensor with a large dynamic range and 2pFLIM enables robust quantitative monitoring of the PTEN conformational state, which serves as a proxy for PTEN activity in complex biological environments.

Following validation and optimization (**Fig. 1, Extended Fig. 1**), we used this approach to examine how PTEN conformation is affected by a number of pathogenic point mutations. The results indicate that certain mutations dramatically affect the PTEN conformation and may produce a destabilized open state. We confirmed these results using structural modeling and can identify a close correlation between FRET measurements and predicted structural stability (**Fig. 2**). While all of the mutations were previously associated with a LOF of PTEN catalytic activity ^17^, our approach uncovers a new link between PTEN dysfunction and structural destabilization. We hypothesize that a pathogenic mutation would lead to reduction in PTEN sensitivity to ongoing changes in signaling and phosphorylation. Future use of this experimental approach will allow us to characterize novel patient specific PTEN mutations as well as enable high-throughput drug screens for PTEN related therapeutics.

One crucial factor to consider in the design of new biosensors is the importance of minimizing their interference with endogenous signaling pathways while preserving normal cellular structure and function. Since our biosensor utilizes a full length PTEN protein, it was important to prevent the previously described interference with endogenous PTEN signaling pathways ^14,31^. Having screened several stable LOF point mutations ^17^, the R14G mutation was selected because of retained conformational changes and the dynamic range of the biosensor. This construct minimally perturbs endogenous PTEN signaling as evidenced by cellular morphology, stability, and development (**Fig. 3**). Importantly, we can demonstrate that long term (weeks) neuronal overexpression of the PTEN biosensor does not affect the spontaneous neuronal activity (**Fig. 6i**). This provides further validation for the utility of our approach and its applicability to study PTEN signaling in the brain with minimal physiological perturbation.

Dynamic imaging of neuronal PTEN activity was conducted in various applications including the well-studied model system *C. elegans*. 2pFLIM of intact worms expressing the sensor enabled us to monitor PTEN neuronal activity at cellular resolution across the lifespan of the intact worm (**Fig. 4**). PTEN is well conserved across species, and the worm homolog, *daf-18,* has been shown to be critical for the metabolic state of the worm and longevity ^53^. Interestingly, the results reveal increases in neuronal PTEN activity that parallel the stages of worm development. Knockout of *daf-2*, a conserved and well-studied IGF1R worm homolog that has a critical role in regulation of proteostasis and longevity ^57^, upstream of PTEN function, resulted in a sustained increase in PTEN activity throughout development (**Fig. 4**). This suggests the presence of a novel direct interplay between PTEN and *daf2* signaling. Previous studies have shown that PTEN overexpression is associated with an increased lifespan in mice ^32^, which was explained by reductions in cellular and tissue proliferation ^14,33^. Future studies will allow to further study developmental and longevity related pathways by monitoring PTEN activity in vivo in small model systems such as *C elegans*.

In vivo 2pFLIM of PTEN activity provides unprecedented spatial resolution of PTEN activity in intact neuronal circuits in the mouse brain where we monitored the neuronal cell body as well as micron sized dendritic spines, the site of synaptic communications (**Fig. 5**). In addition, we used CRISPR/Cas9 to manipulate IGF1R and Tsc2, which are upstream and downstream of PTEN, respectively, in order to examine how different perturbations in the signaling pathway impact neuronal PTEN activity. Genetic perturbation of IGF1R and Tsc2 led to similar increases in phosphorylation of S6 (pS6), a well validated indicator of PTEN signaling. Interestingly, we found that each manipulation led to distinct compartmentalization of PTEN signaling, which correlated with distinct synaptic changes; while somatic PTEN activity was upregulated and associated with larger cell size, dendritic PTEN activity was affected differently after KO of Tsc2 compared to KO of IGF1R, reflecting distinct morphological alterations. Future studies will be needed to determine whether synaptic dysregulation of compartmentalized PTEN activity is a conserved feature following various perturbations of synaptic structure and function.

To study how PTEN signaling is regulated in respect to other signaling pathways or neuronal activity, we chose to engineer a red-shifted biosensor variant, R-PTEN (**Fig. 6**). This version relies on the previously established red-shifted FRET/FLIM pair, mCyRFP2-mMaroon1 ^37,42^. The advantage of using a red-shifted FRET pair is that there is a wide variety of biosensors that employ GFP based fluorescence. Our choice of label permits simultaneous imaging of PTEN alongside a large repertoire of GFP based biosensors. In addition, this combination is particularly suitable for in vivo imaging since the use of dual lasers for excitation is technically challenging. The combination of R-PTEN and the sensitive calcium sensor GCaMP8 allows simultaneous monitoring of PTEN activity and ongoing neuronal activity in single cells, in awake animals. Interestingly, the results reveal a negative correlation between PTEN activity and spontaneous neuronal activity. Previous studies have reported significant changes in neuronal activity and hyperexcitability after perturbing PTEN. Our results pave the way for future studies using this approach, and further unravel the dynamics of the interplay between PTEN and neuronal function in the brain.

Lastly, we combined genetic targeting and spectral PTEN sensor variants to explore a critical and unresolved question: Do PTEN signaling dynamics differ between cell types in the brain? Previous studies addressed this question by cell-type specific genetic perturbation of PTEN, which is contentious since PTEN KO leads to drastic morphological and functional dysregulation in all cell-types tested so far. It was therefore difficult to determine whether the signaling dynamics of PTEN are regulated in cell-specific manner or similar in all cells. Our newly developed biosensor now provides evidence for distinct regulation of PTEN activity in excitatory and inhibitory cells in the cerebral cortex (**Fig. 7**). We found that sensory deprivation led to differential changes in PTEN activity in inhibitory and excitatory cells. Previous work suggested that PTEN activity in inhibitory cells is critical for the establishment of E/I balance in the cortex ^18^. This is a vital feature impacted by neurodevelopment disorders such as autism spectrum disorders ^68^, which are closely associated with PTEN mutations ^24^. Future studies using the PTEN biosensor will allow us to analyze the cell-specific contribution of PTEN to the development of neuronal circuits and will shed light on the emergence of ASD and related neurodevelopmental disorders.

In summary, we have engineered a novel FRET based PTEN biosensor, which is optimized for 2pFLIM. This new tool allows us to explore the spatial and temporal dynamics of PTEN in a broad range of *in vitro and in vivo* biological systems and identify the contribution of cell-specific PTEN activity to cellular structure and function.

## Materials and Methods

### Animals

Animal experiments were approved by the Institutional Animal Care and Use Committee in Tel Aviv University. In Utero Electroporation (IUE) was performed on time pregnant ICR mice (Envigo). For AAV injections, ICR females were crossed with Homozygous PV Cre male mice (Stock No. 017320, The Jackson Laboratory). Heterozygous litters were used for AAV injection in the first postnatal week. Both male and female mice were used throughout experiments. All animals were kept in a normal light/dark cycle (12 h/12 h, lights on at 7AM) and had free access to food and water.

### Plasmids and AAV construction

For construction of the G-PTEN sensor, mEGFP followed by a linker of SGLRSA were fused to the N terminus of the PTEN coding sequence (a gift from Jose Esteban, Addgene plasmid # 110181). On the C terminus of PTEN, a linker of PTP was followed by coding sequence for sREACh. For the R-PTEN version, the donor and acceptor were replaced by mCyRFP2 ^37^ and mMaroon1 ^69^, respectively. PTEN point mutations described in the results and the 4A mutant were generated using PCR and the Q5 mutagenesis kit (NEB). RhoA plasmids were previously described in ^42^. For experiments in cell lines, The PTEN sensor was used under the CMV promoter (EFPC-C1 backbone). For In Utero Electroporation (IUE) in mice, the constructs were cloned under the CAG promoter. For co-expression in astrocytes and in neurons using IUE, the piggyBac (PB) transposon system was used together with expression pCAG-hyPBase ^70^. For production of AAVs, we replaced the acceptors for sREACh and mMaroon for G and R versions with a codon-diversified versions ^61^ to avoid AAV mediated cleavage due to high nucleotide homology of donor and acceptor. AAV constructs contained the human Syanpsin promoter or a Cre dependent FLOXed version under the CAG promoter ^42^. For expression in C. Elegans, we used the pan-neuronal IRP-I7 promoter ^54^. The G-PTEN FLOXed AAV was packaged by the ETH viral core and was a gift by Dr. Aiman Samir Saab in PHP serotype, the Synapsin R-PTEN sensor were packaged in AAV serotype 9 by the ELSC vector core facility. AAVs containing GCaMP8s under the Synapsin promoter were purchased from Addgene (Plasmid #162374).

For gene editing via CRISPR/Cas9, we used the PX330 backbone (a gift from Feng Zhang, Addgene plasmid 42230). We introduced gRNA sequences according to target gene (5’-3’): PTEN (TCACCTGGATTACAGACCCG, exon 6), Tsc2 (TGTTGGGATTGGGAACATCG, exon 2), IGF1R (GAAAACTGCACGGTGATCGA, exon 2), and a scrambled control based on PTEN exon 6 (ACTCGAGACGCGCATCTACT). The gRNA Sequence for targeting PTEN was generously provided by Dr. Brian Luikart. The other gRNA sequences were identified in the second exon, by choosing the highest scoring sequence for sensitivity and specificity ^71^.

Plasmids were constructed using standard molecular biology methods including Polymerase Chain Reaction, Gibson Assembly, Enzyme Restriction Reactions and Q5 Side Directed Mutagenesis (NEB). For optimization, linkers were edited using PCR, and mutations were introduced using PCR. All products were verified using Sanger Sequencing.

### Cell culture maintenance

HEK293T cells (ATCC) in passage number 12-20 were cultured in DMEM supplemented with 10% FBS, 1% L-Glutamic Acid and 1% Penicillin-Streptomycin at 37°C in 5% CO2 and transfected with plasmids (Mirus TransIT-LT1 Transfection Reagent). Imaging was performed 24hr following transfection in external Tyrode solution (119mM NaCl, 5mM KCl, 25mM HEPES, 2mM CaCl2, 2mM MgCl2, 33mM Glucose) at pH 7.4. TBB (Tocris) was administered with a concentration of 50μM in 2ml. For washout experiments, HEK cells were grown on PLL coated slides to increase adhesion, the TBB solution was removed 1hr later from the plate and replaced with 2ml of Tyrode with or without 100ng/ml EGF (PeproTech) for an additional 2hr. For RhoA co-expression with PTEN sensor experiments, plasmids were used in a 1:1 ratio. HEK Cells were used as an expression and experimentation platform only and were not rigorously tested for potential contamination from other cell lines.

### In silico mutation analysis

To evaluate the effect of mutations, the latter were incorporated within the available PTEN structure (PDB: 1D5R) via the Maestro BioLuminate suite in the Schrodinger software. The stability change (ΔG) of the protein was calculated based on the Prime energy function with an implicit solvent term and is defined as the free energy difference between the folded and the unfolded state. Stability relative to the wild-type protein (ΔΔG) was calculated based on the residue scanning calculation module. A negative value points to a more stable mutant.

### C elegans transgenic worms

The GFP-PTEN-sREACh R14G was cloned under a Psnb*-1* pan-neuronal ^54^. For *C. elegans* transgenesis, 50 ng of plasmid was injected into the gonad along with 50 ng of a P*myo-2*::dsRed pharyngeal marker. The strain was maintained on nematode growth medium agar seeded with OP50 Escherichia coli at 20°C. Before experiments, worms were synchronized using bleach and the next generation was used for each experiment.

For TBB experiments, 3 days after bleach synchronization 30 fluorescent positive worms at L4 developmental stage were picked onto TBB plates. TBB plates were made by mixing 100μl M9 buffer and 500μM TBB, seeding the solution on the OP50 E. Coli plates, and letting dry at 20-25°C for 1hr in sterile conditions before bleach synchronizing *C. elegans* to the plate. 72 hours later, fluorescent positive adult *C. elegans* were mounted onto a slide with 30μl M9 buffer and 3μl Levamisole at 1μM. Images were taken of neurons with Tau1 fixed at 3.2ns and Tau2 fixed at 1.6ns. The same routine was done for the control group, without adding TBB to the plates.

For *daf-2* RNAi experiments worms were fed HT115 (DE3) bacteria expressing the L4440 vector containing *daf-2* specific sequence. Empty L4440 plasmid was used as the negative control. RNAi plates were prepared with NGM media containing 10mM IPTG and 100 μg/ml of ampicillin. For dsRNA induction, an overnight culture of bacteria was diluted 1:50 and grown at 37°C with shaking for 3-4 hours, after which IPTG was added to a final concentration of 10mM and allowed to grow for an additional 3-4 hrs. This culture was used to seed the RNAi plates, air dry and grow overnight at room temperature. Gravid hermaphrodites were bleached on the RNAi plates to release the embryos. 72 hours later, fluorescent positive worms were morphologically sorted by larval stage (L1, L2/3, L4, Adult) and mounted for imaging with 30μl M9 buffer and 3μl Levamisole at 1μM. Images were taken of neurons with Tau1 fixed at 3.2ns and Tau2 fixed at 1.6ns.

### Immunohistochemistry

Perfused brains of mice expressing the tested gRNA were sliced at 100μm using a vibratome (Leica VT1000S) and thereafter stained. Slices were washed three times with PBS for 5 minutes each at room temperature and incubated in PBST (1.2% triton) for 15 minutes. Slices were washed an additional 3 times for 5 minutes each and transferred for 1 hour in blocking buffer (5% NGS, 2% BSA, 0.2% Triton in PBS solution) at room temperature. The slices were then incubated in primary rabbit anti-phospho-S6 S235/236 1:200 diluted in blocking buffer, (Cell Signaling#4858) overnight at 4°C. Slices were then washed three times with PBS for 5 minutes and incubated for 90 minutes in secondary Alexa 594 1:1000 (Thermo Fisher Scientific). Slices are washed three times with PBS for 5 minutes each, mounted onto slides, sealed with mounting medium (VectaShield HardSet) and left to dry overnight.

### Confocal microscopy

Images were taken using a ZEISS LSM 900 in confocal mode, using Zen 3.1. Fluorescence readouts were obtained at ex/em of 488/507 nm, 594/614 nm for green and red, respectively.

### In Utero Electroporation, AAV injections and cranial windows

In Utero Electroporation was performed as previously reported ^48^. Briefly, ICR E14.5-15.5 timed pregnant mice (Envigo) were anesthetized with isoflurane and given 0.1 mg buprenorphine SR for analgesia. Uterine horns were exposed though an abdominal incision along the Linea Alba, and the right lateral ventricle of each embryo was injected with plasmids, at a final concentration of 1 μg/μl with 0.01% Fast Green dye (Sigma-Aldrich). Five electrical pulses (40 V, 50-ms duration, 1 Hz) were delivered using a NEPA21 electroporator (NEPAGENE) with a triple electrode configuration. After birth, fluorescence in the cortex of the pups were visually validated under a fluorescent lamp. G-PTEN was used at a concentration of 1.5µg/µl. For injections using PX330 – we used PX330, CAG-CyRFP and PTEN plasmids at final concentration of 1µg/µl each.

For AAV injections, we used a mix of Syn R-PTEN and FLEX_G-PTEN (2×10^13^/2×10^13^ vg/ml) or R-PTEN and GCaMP8 (2×10^13^/1×10^13^ vg/mL). Mouse pups (p3-p4) were anesthetized with isoflurane and the right somatosensory cortex was targeted using a glass pipette with total of 1µl AAV. Pups were then left to recover on a heating pad with their litter and following recovery were returned to their home cage until weaning.

For cranial window surgery, mice (p21-70) were deeply anesthetized with isoflurane for induction (2%–3%), and administered 1 μg/g buprenorphine SR for analgesia, and 5 mg/kg carprofen to prevent edema and inflammation. Following fixation in a stereotaxic frame, hair was removed and skin and skull exposure. Then, a 2.5-3mm circular craniotomy was performed over the imaging site using a dental drill. We verified positive fluorescence from IUE/AAV at the somatosensory cortex. The skull was sealed using a 3mm, No.1 circular cover glass glued on a 5mm circular cover glass and cemented to skull along with a head plate to secure the head during imaging using dental cement (C and B Metabond, Parkell). Mice were left to recover and were used for in vivo imaging according to experimental design. Mice that showed signs of window occlusion, tissue damage were excluded from imaging and analysis.

### 2pFLIM microscopy

We use a 2pFLIM microscope which was based on a Galvo-Galvo two-photon system (Thorlabs) and a 2pFLIM module (Florida Lifetime Imaging), equipped with a Time-Correlated Single Photon Counting board (Time Harp 260, Picoquant). The microscope was controlled via the FLIMage software. For excitation, we used a Ti:Sapphire laser (Chameleon, Coherent) at a wavelength of 920 nm. Excitation power was adjusted using a pockel cell (Conoptics) to 1.0–2.0 mW for in vitro experiments and 5-40mW for in vivo experiments. Emission was collected with a 16 × 0.8 NA objective (Nikon), divided with a 565-nm dichroic mirror (Chroma), with emission filters of 525/50 nm and 607/70 nm, detected with two Photo-Multiplier Tubes with low transfer time spread (H7422-40p, Hamamatsu). Images were collected by 128 × 128 or 256 x 256 pixels and acquired at 2 ms/line, averaged over 24 frames. In vivo imaging of R-PTEN/GCaMP was collected at frame rate of 4Hz and analyzed offline.

### 2pFLIM analysis

All FLIM analysis was performed using a custom C# software (available on the Yasuda lab Github: https://github.com/ryoheiyasuda/FLIMage_public). Fluorescence lifetime decay curve A(t) was fitted with a double exponential function convolved with the Gaussian pulse response function:

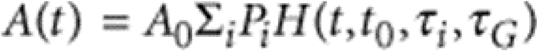

Pi- is the fractional population with the decay time constant of τi, and A0 is the initial fluorescence before convolution. H(t) is an exponential function convolved with the Gaussian instrument response function (IRF).

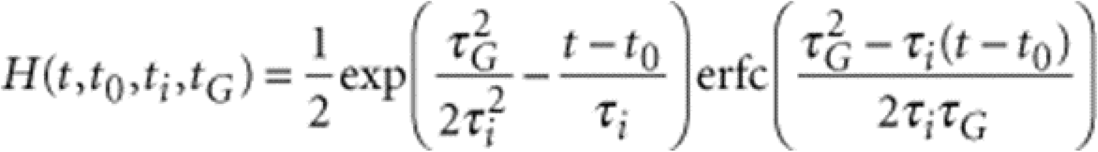

erfc is the error function, τG is the width of the Gaussian pulse response function, and t0 is the time offset. Weighted residuals were calculated using:

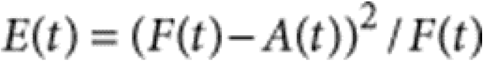

Fitting was performed by minimizing the summed error δ2 = ΣtE(t) for parameters t0, τi (i = 1,2) and τG. We created fluorescence lifetime images by finding the averaged fluorescence lifetime (τm) by the mean photon arrival time subtracted by t0 in each pixel as:

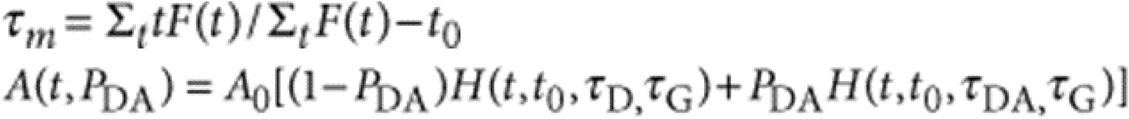

where t0 and τG are obtained from a curve fitting to the fluorescence lifetime decay averaged over all ROIs in an image.

The fluorescence lifetime of mEGFP / mCyRFP was fixed for TauD/DA values of 2.6/1.1, 3.2/1.7, and 3.65/1.4 ns for G-PTEN in mammalian G-PTEN, *C elegans* G-PTEN, and R-PTEN respectively. Mean lifetimes were sampled from ROIs, as well as their total photon numbers for intensity. We found no correlation between lifetime and fluorescence intensity for the PTEN sensor.

### GCaMP analysis

Movie registration and signal extraction were processed using a custom Python script written by G. Bond. XY motion correction was processed and exported using CaImAn (FlatIronInstitute). ROIs were drawn using ImageJ’s Cell Magic Wand plugin. In each movie, changes in fluorescence (ΔF) were computed relative to the baseline fluorescence (F0) to produce ΔF/F. The cumulative sum of ΔF/F was calculated by setting negative values as 0.

### Statistical analysis

All values are presented as mean ± s.e.m. Statistical significance was tested by two-tailed T test for comparison of two groups, or one way ANOVA followed by post-hoc Tukey’s multiple comparison test for comparison of multiple groups (P < 0.05) using GraphPad Prism 9.0 (GraphPad Software). For correlation analysis, outliers were removed using the Grubbs method with an Alpha of 0.01. For correlation analysis, we used two-tailed Pearson test followed by simple linear regression for graphic representation. For differences in variability, the F test was used (P < 0.05). For all statistical tests * = p < 0.05, ** = p < 0.01, *** = p < 0.001 and **** = p < 0.0001 were considered significant. Sample sizes were not predetermined using statistical methods.

## Data and Code Availability

The data supporting the current study are available upon request. Software for analysis of the FLIM data is available on the Ryohei Yasuda lab Github: https://github.com/ryoheiyasuda/FLIMage_public.

Plasmids from this paper will be deposited in Addgene.

## Acknowledgments

We would like to thank Long Yan for help with FLIM microscopy setup, Brian Luikart for generously providing the gRNA sequence for PTEN, Ithai Rabinowitch for providing the neuronal *C. elegans* promoter, Ehud Cohen for the Daf2 RNAi, and Inna Slutsky for PV-Cre mouse. AAVs for R-PTEN were produced by the ELSC viral facility in the Hebrew University. Aiman S. Saab and the University of Zurich viral facility generously provided custom AAVs for FLEX-G-PTEN. Greg Bond and Ben Scholl for assistance with GCaMP analysis, and Laviv lab members for discussions. This work was supported by a grant from the Autism Research Institute (ARI), T.L. is supported by a Ben Barres Early Career award from the Chan Zuckerberg Initiative (CZI), by the Israel Science Foundation (ISF) grants 1384/21 and 1385/21, and an ERC starting grant number 101040128. TK is supported by a Teva BioInnovators fellowship. M. Gabay is supported by a PhD scholarship from the Yoran Institute for Human Genome Research and the Gertner Institute for Medical Nanosystems at Tel Aviv University.

## Author Contributions

T.L. conceived the project. T.K. and T.L. performed all in vitro and in vivo preparations and experiments, analysis and writing. M. Gabay and M. Gal performed computational prediction. Y.L. assisted with plasmid preparations. S.E assisted with immunostaining. N.M. and M.M assisted in IUE. A.N. and R.Z.B prepared and advised on the PTEN C. elegans model. T.L. and T.K. wrote the paper with feedback from all the authors.

