## Supplementary Figures for "Genetically encoded biosensor for fluorescence lifetime imaging of PTEN dynamics in the intact brain"

Extended Figure 1

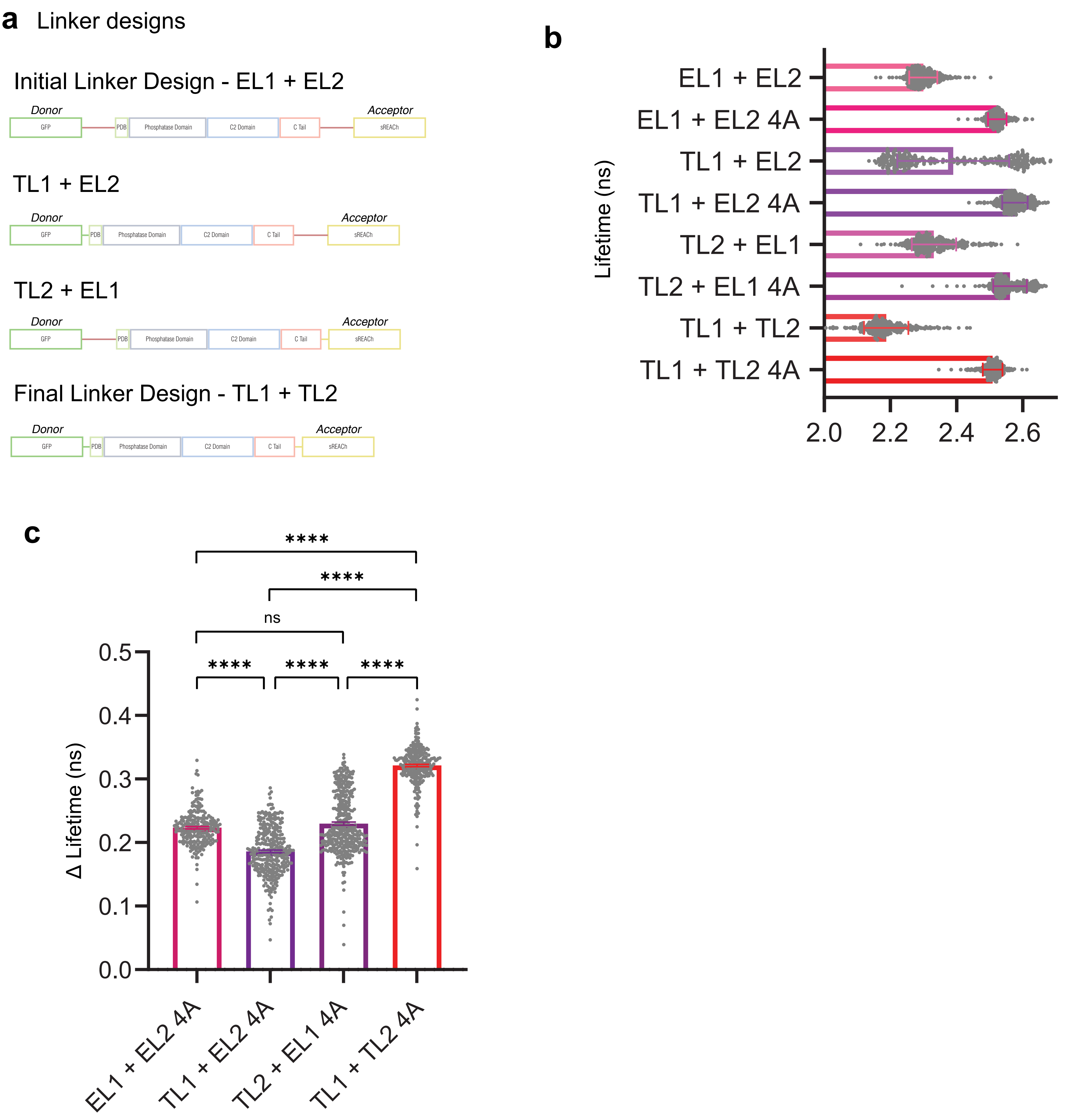

#### Extended Figures legends:

##### Extended Figure 1

(a) Schematic design of the different FRET/FLIM-based mEGFP-PTEN-sREACh sensor linker versions; combinations of extended linkers on either side of mEGFP-PTEN-sREACh (EL1, EL2 respectively) and truncated linkers on either side of mEGFP-PTEN-sREACh (TL1, TL2 respectively).

(b) Quantification of fluorescence lifetime of the different mEGFP-PTEN-sREACh linker versions. HEK293 were transfected with either EL1 + EL2 (2.30±0.002 ns, n = 310), EL1 + EL2 4A (****, 2.52±0.002 ns, n = 223), TL1 + EL2 (2.39±0.009 ns, n = 320), TL1 + EL2 4A (****, 2.58±0.002 ns, n = 310), TL2 + EL1 (2.33±0.004, n = 320), TL2 + EL1 4A (****, 2.56±0.002 ns, n = 400), TL1 + TL2 (2.19±0.004 ns, n = 288) or TL1 + TL2 4A (****, 2.51±0.002 ns, n = 281). Comparisons between WT and 4A mutation of each variant.

(c) Quantification of change in mean fluorescent lifetime (Δ) of the different mEGFP-PTEN-sREACh linker version 4A mutations from WT; EL1 + EL2 4A (0.22±0.002 ns, n = 223), TL1 + EL2 4A (0.19±0.002 ns, n = 310), TL2 + EL1 4A (0.23±0.002 ns, n = 400), TL1 + TL2 4A (0.32ns ±0.002 ns, n = 281). ns $p = 0.2106$.

Error bars represent s.e.m; statistical differences were measured using one-way ANOVA followed by post-hoc Tukey’s multiple comparison test. **** $p < 0.0001$.

Extended Figure 2

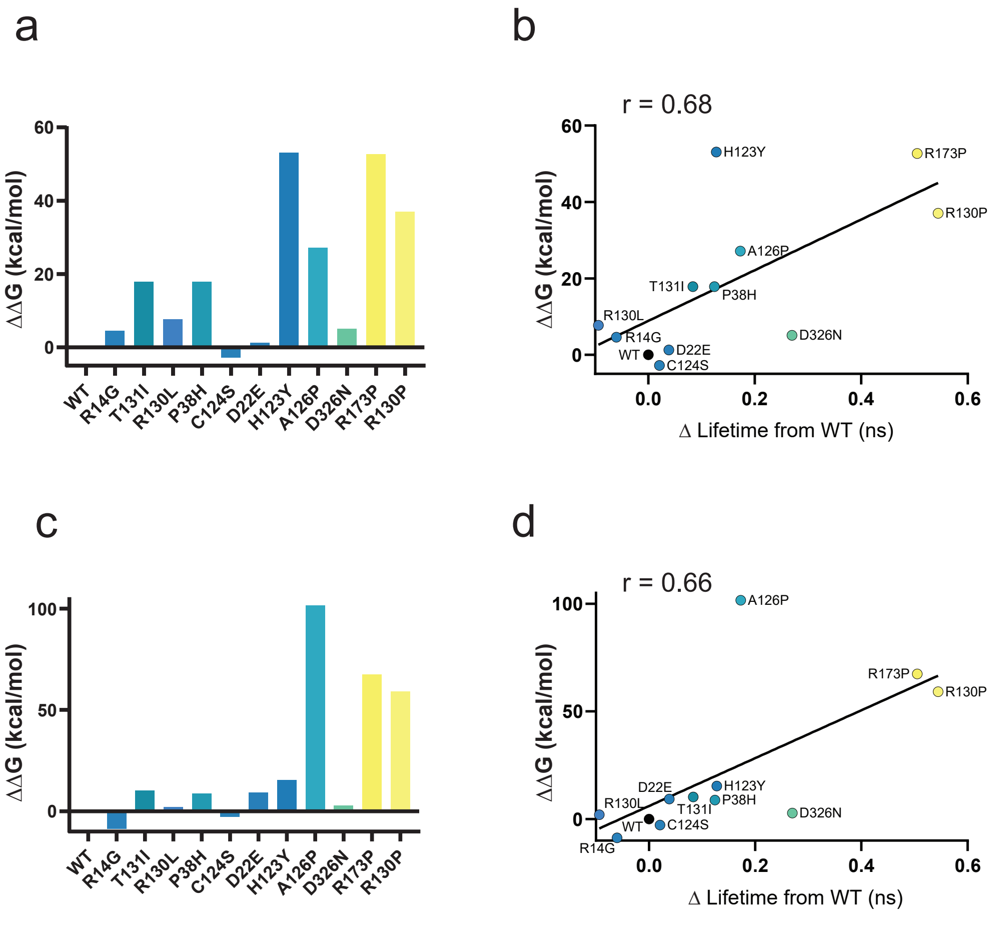

**Extended Figure 2**

(a) Predicted $\Delta\Delta$G resulting from changes in electrostatic interactions of the mEGFP-PTEN-sREACh mutants compared to WT. R14G (4.58), T131I (17.82), R130L (7.7), P38H (17.87), C124S (-2.77), D22E (1.29), H123Y (53.06), A126P (27.17), D326N (5.08), R173P (52.7), R130P (37.03).

(b) Correlation between $\Delta\Delta$G of PTEN mutants and their corresponding measured changes in fluorescence lifetime of mEGFP-PTEN-sREACh mutants compared to WT mean ($p = 0.015$, $r = 0.68$).

(c) Predicted $\Delta\Delta$G resulting from changes in covalent interactions of the mEGFP-PTEN-sREACh mutants compared to WT. R14G (-8.75), T131I (10.27), R130L (2.07), P38H (8.83), C124S (-2.77), D22E (9.27), H123Y (15.37), A126P (101.64), D326N (2.82), R173P (67.38), R130P (59.12).

(d) Predicted $\Delta\Delta$G resulting from changes in covalent interactions of the mEGFP-PTEN-sREACh mutants compared to WT. mean ($p = 0.020$, $r = 0.66$).

Correlation was measured using two-tailed Pearson test and plotted using simple linear regression. One computer predicted outlier (G127R) was found and removed using Grubbs test for outliers (Alpha = 0.01).

Extended Figure 3

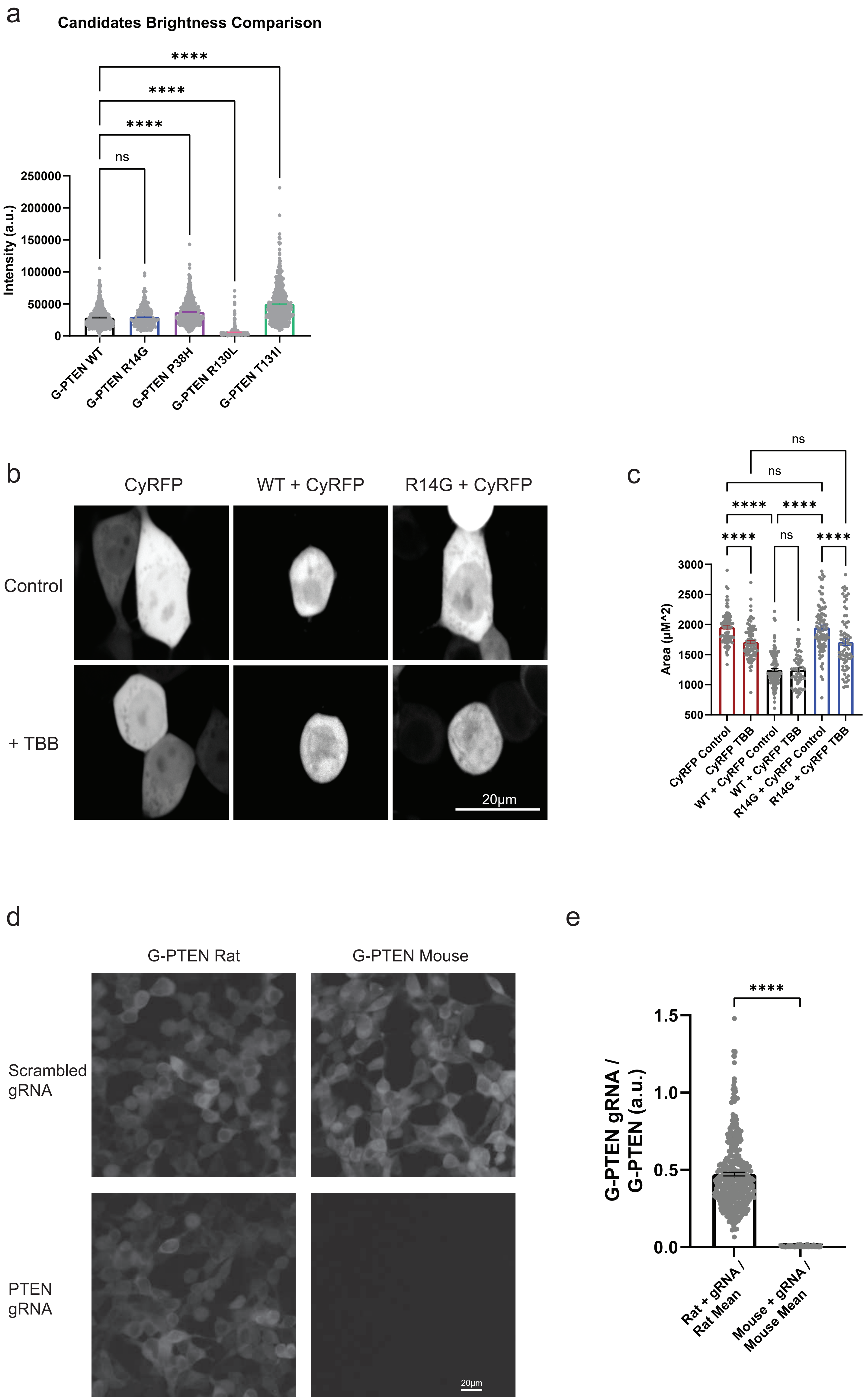

**Extended Figure 3**

(a) Quantification of fluorescent intensity in HEK293 cells expressing mEGFP-PTEN-sREACh WT (28503±670 units, n = 504), R14G (29750±1105 units, n = 200), P38H (37229±859 units, n = 480), R130L (5788±441 units, n = 353) or T131I (49850±1292 units, n = 495). ns p = 0.9396.

(b) Representative images of PTEN sensor fluorescent intensity before and after 3hr of TBB (50 µM). HEK293 cells were co-transfected with either CyRFP alone, co-expressing mEGFP-PTEN-sREACh WT, or mEGFP-PTEN-sREACh R14G. Scale bar; 20 µm.

(c) Quantification of cell area of CyRFP alone, before (1958±33.40 $\mu m^2$, n = 82, 1709±30.15 $\mu m^2$, n = 88), + mEGFP-PTEN-sREACh WT (1245±26.78 $\mu m^2$, n = 109, 1247±34.35 $\mu m^2$, n = 62), or with mEGFP-PTEN-sREACh R14G (1951±42.25 $\mu m^2$, n = 92, 1707±53.80 $\mu m^2$, n = 75) before and after TBB application, respectively. ns p > 0.9999 for CyRFP Control compared to R14G + CyRFP Control, for CyRFP TBB compared to R14G + CyRFP TBB, and for WT + CyRFP Control compared to WT + CyRFP TBB.

(d) Representative images of G-PTEN fluorescence intensity with or without co-expressing spCas9 and PTEN gRNA. G-PTEN was modified from rat sequence which differs from gRNA in one nucleotide, to mouse PTEN which matches the gRNA template. Scale bar; 20 µm.

(e) Quantification of sum fluorescent intensity of the PTEN biosensor co-expressing PTEN gRNA with the mean sum of G-PTEN alone, for the rat based version (0.473 ± 0.01, n = 380) and mouse based sensor (0.003 ± 0, n = 400).

Error bars represent s.e.m; statistical differences for (a, c) were measured using one-way ANOVA followed by post-hoc Tukey's multiple comparison test. Statistical difference for (e) was measured using unpaired two-tailed student t-test. **** p < 0.0001.

Extended Figure 4

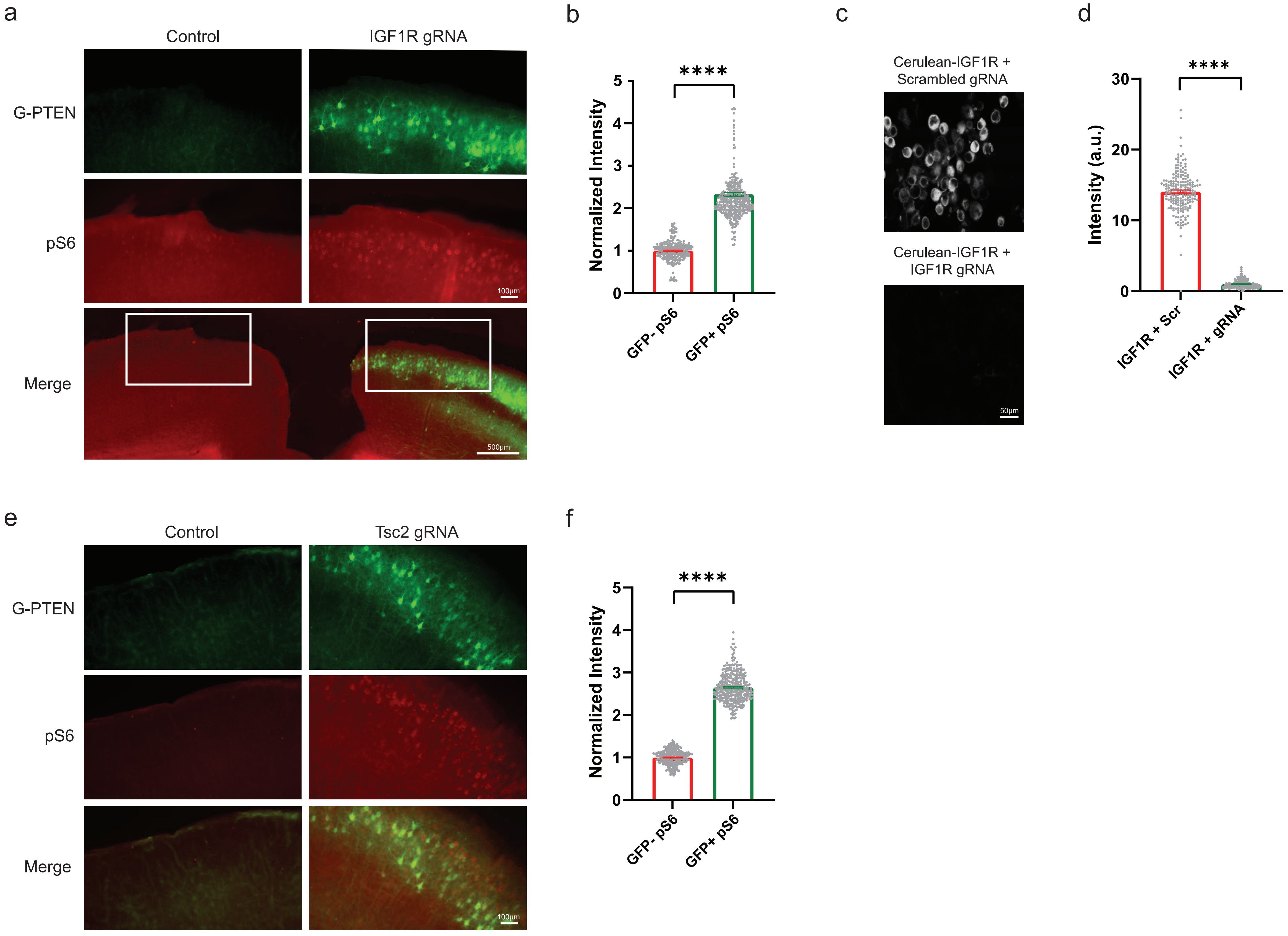

**Extended Figure 4**

(a) Representative images of pS6 stained cortical brain slices, in WT hemisphere (control) and the hemisphere expressing G-PTEN, spCas9 and an IGF1R targeted gRNA. Zoomed scale bar; 100 µm. Wide field scale bar; 500 µm.

(b) Quantification of pS6 normalized intensity on stained cortical brain slices, within the WT hemisphere not expressing GFP (1±0.145 units, n = 261) and gRNA targeted IGF1R hemisphere co-expressing GFP (2.33±0.044 units, n = 349).

(c) Representative images of HEK293 co-expressing Cerulean tagged IGF1R, spCas9 and either scrambled control gRNA or IGF1R targeted gRNA. Scale bar; 50 µm.

(d) Quantification of Cerulean-IGF1R normalized intensity in HEK293 cells with either scrambled (Scr) control gRNA (14.07±0.205 units, n = 201) or IGF1R targeted gRNA (1±0.039 units, n = 212).

(e) Representative images of pS6 stained cortical brain slices, in WT hemisphere (control) and the hemisphere expressing G-PTEN, spCas9 and an Tsc2 targeted gRNA. Scale bar; 100 µm.

(f) Quantification of pS6 normalized intensity on stained cortical brain slices, within the WT hemisphere not expressing GFP (1±0.009 units, n = 340) and gRNA targeted hemisphere co-expressing GFP (2.65±0.019 units, n = 342).

Error bars represent s.e.m; statistical difference for (b, d, f) were measured using unpaired two-tailed student t-test. **** $p < 0.0001$.

### Extended Figure 5

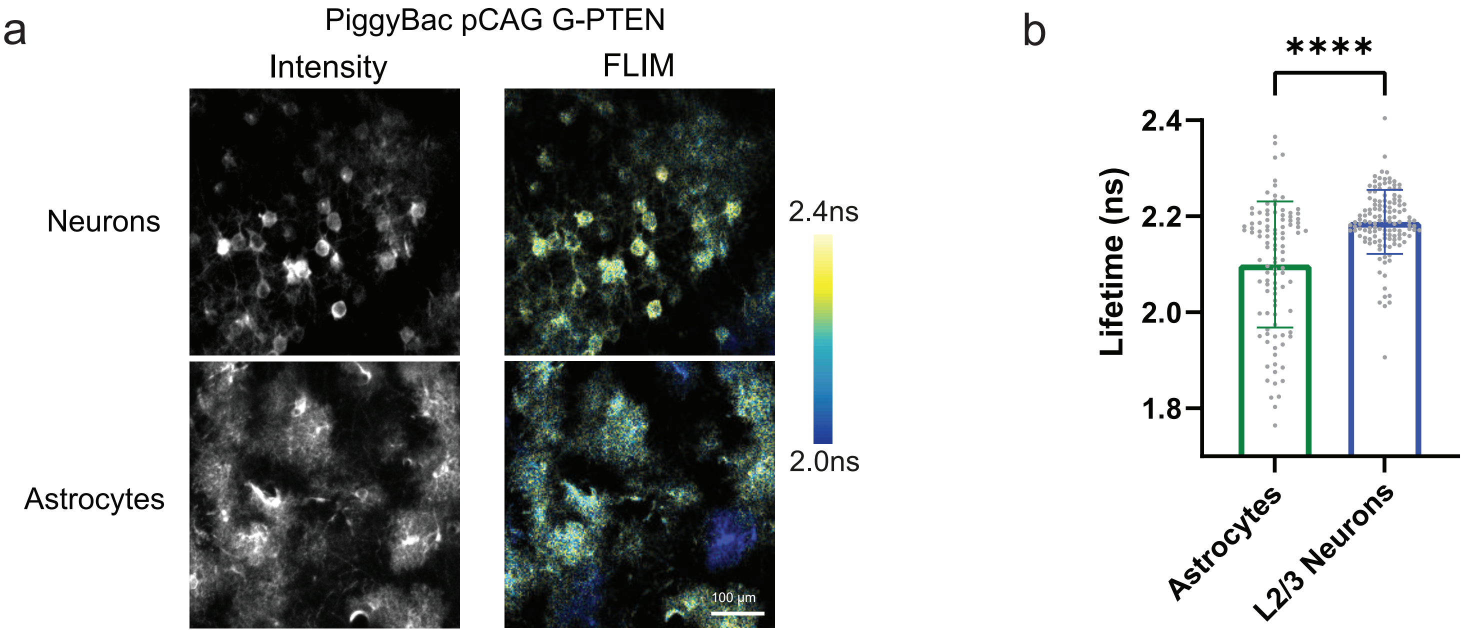

**Extended Figure 5**

(a) Representative images of fluorescence intensity and FLIM of L2/3 neurons and astrocytes expressing G-PTEN after PiggyBac IUE injection. Scale bar; 100 µm.

(b) Quantification of fluorescence lifetime in astrocytes (2.09±0.013 ns, $n = 104$) and in L2/3 neurons (2.19±0.006 ns, $n = 129$) in the same mice.

Error bars represent SEM; statistical differences for (b) were measured using one-way ANOVA followed by post-hoc Tukey's multiple comparison test. **** $p < 0.0001$.

Extended Figure 6

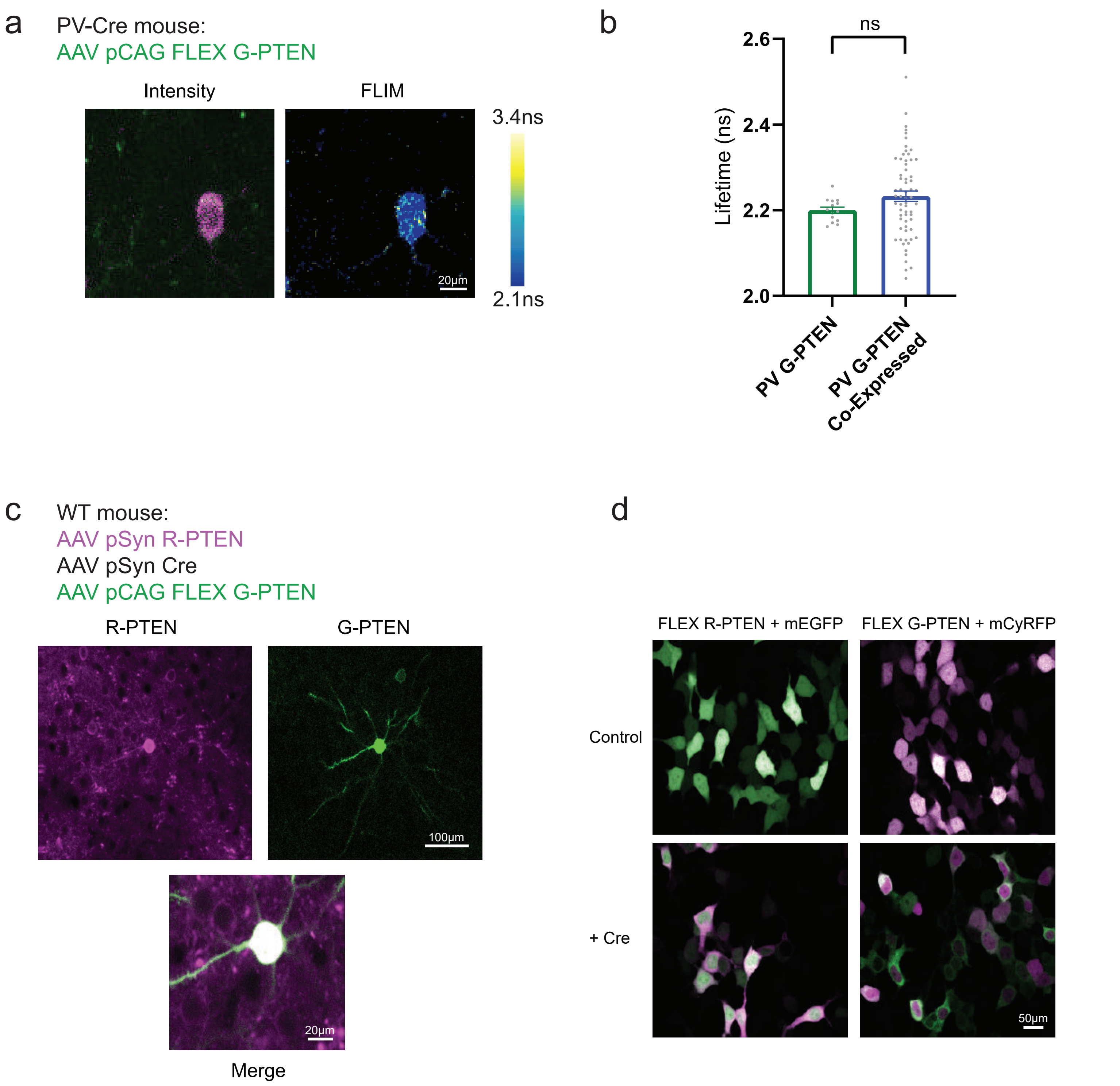

**Extended Figure 6**

(a) Representative fluorescence intensity and lifetime images of PV-Cre mouse after AAV injection of Cre dependent G-PTEN. Scale bar; 20 µm.

(b) Quantification of fluorescence lifetime in mice with PV targeted G-PTEN (2.20±0.007ns, n = 14) and in mice co-expressing G-PTEN in PV cells and R-PTEN expressing in excitatory neurons. (2.23±0.012ns, n = 64). ns $p = 0.2165$.

(c) Representative 2p fluorescence intensity images of WT mouse after simultaneous AAV injection of Syn R-PTEN, Syn Cre and FLEX G-PTEN. Widefield scale bar; 100 µm. Zoomed scale bar; 20 µm.

(d) Representative images of HEK293 without or with Cre, co-expressing the FLEX R-PTEN or FLEX G-PTEN sensors with mEGFP or mCyRFP markers, respectively. Scale bar; 50 µm.

Error bars represent SEM; statistical difference for (b) was measured using unpaired two-tailed student t-test.

**Movie 1 – Dynamics of G-PTEN activity following TBB application in HEK293**

A movie showing changes in PTEN lifetime following TBB application. Images taken every 10 minutes following over 200 minutes. Frame dimensions; 195 µm x 195 µm.

**Movie 2 – Simultaneous in vivo imaging of GCaMP8s and R-PTEN in the mouse brain**

*In vivo* 2pFLIM imaging of L2/3 cells in the somatosensory cortex co-expressing GCaMP8s (green) and R-PTEN (magenta). Movie is displayed at 16 frames per second, acquired at 4 frames per second. Frame dimensions; 195 µm x 195 µm.
